# Discovering Novel Circuit Mechanisms in Higher Cognition through Factor-Centric Recurrent Neural Network Modeling

**DOI:** 10.64898/2026.04.16.718933

**Authors:** Yiteng Zhang, Xingyu Li, Xuewen Shen, Fangting Li, Gouki Okazawa, Liping Wang, Jianfeng Feng, Bin Min

**Affiliations:** Lingang Laboratory, Shanghai, China; School of Data Science, Fudan University, Shanghai, China; School of Physics, Center for Quantitative Biology, Peking University, Beijing, China; Institute of Neuroscience, Key Laboratory of Brain Cognition and Brain-Inspired Intelligence Technology, CAS Center for Excellence in Brain Science and Intelligence Technology, International Center for Primate Brain Research, Chinese Academy of Sciences, Shanghai, China; Institute of Science and Technology for Brain-Inspired Intelligence, Fudan University, Shanghai, China; Key Laboratory of Computational Neuroscience and Brain-Inspired Intelligence, Fudan University, Ministry of Education, Shanghai, China

## Abstract

Recurrent neural networks (RNNs) have transformed how systems neuroscientists generate hypotheses about circuit mechanisms in higher cognition. Yet their promise has been constrained by a fundamental limitation: conventional RNNs are neuron-centric and therefore often difficult to interpret mechanistically. Here we introduce Restricted-RNN, a factor-centric RNN modeling framework developed to uncover interpretable circuit mechanisms. By formalizing factor communication through subpopulations, the dual of neuron communication through subspaces, Restricted-RNN departs fundamentally from standard neuron-centric RNN models and provides a distinct framework for describing circuit mechanisms. Using this approach, we identify novel circuit mechanisms underlying sequence working memory control and the counterintuitive firing-rate reversal observed in perceptual decision-making, with key predictions supported by neurophysiological recordings from monkey frontal and parietal cortex. More importantly, factor-centric RNN modeling reveals a unified low-dimensional neural control state space that links seemingly disparate phenomena across tasks, providing a geometric framework for understanding the pervasive role of control in higher cognition.

## Introduction

Elucidating circuit mechanisms underlying complex behaviors arguably is one of the major endeavors in neuroscience. The enormous complexity exhibited in single neural activities (Rigotti et al., 2013; Tye et al., 2024), however, poses a major challenge towards this grand endeavor. One promising and influential idea to address this challenge is the computation-through-dynamics framework in which the neural computation underlying behavior is casted as a dynamical process in high-dimensional neural state space, capable of accommodating both single-neuron-level complexity and collective-level simplicity (Churchland et al., 2012; Sussillo, 2014; Vyas et al., 2020; DePasquale et al., 2023; Fig. 1A). This framework is further corroborated with an artificial recurrent neural network (RNN) modeling approach (Fig. 1B), which has been widely adopted to generate new circuit mechanism hypotheses for cognitive and motor tasks (Mante et al., 2013; Sussillo et al., 2015; Rajan et al., 2016; Song et al., 2016; Cueva & Wei, 2018; Wang et al., 2018; Yang et al., 2019; Masse et al., 2019; Whittington et al., 2020; Sorscher et al., 2023; Driscoll et al., 2024; Langdon & Engel, 2025; Li et al., 2025; Voigts et al., 2025). While extremely powerful, this RNN-based modeling approach is of “black-box” nature, oftentimes leading to models difficult to be interpreted, inconsistent with neural data or not comprising the full set of biological solutions (Tolmachev & Engel, 2025; Pagan et al., 2025). This limitation hinders its applications in generating circuit mechanism hypotheses for challenging problems, calling for a major revision or an alternative with higher interpretability.

**Figure 1.**
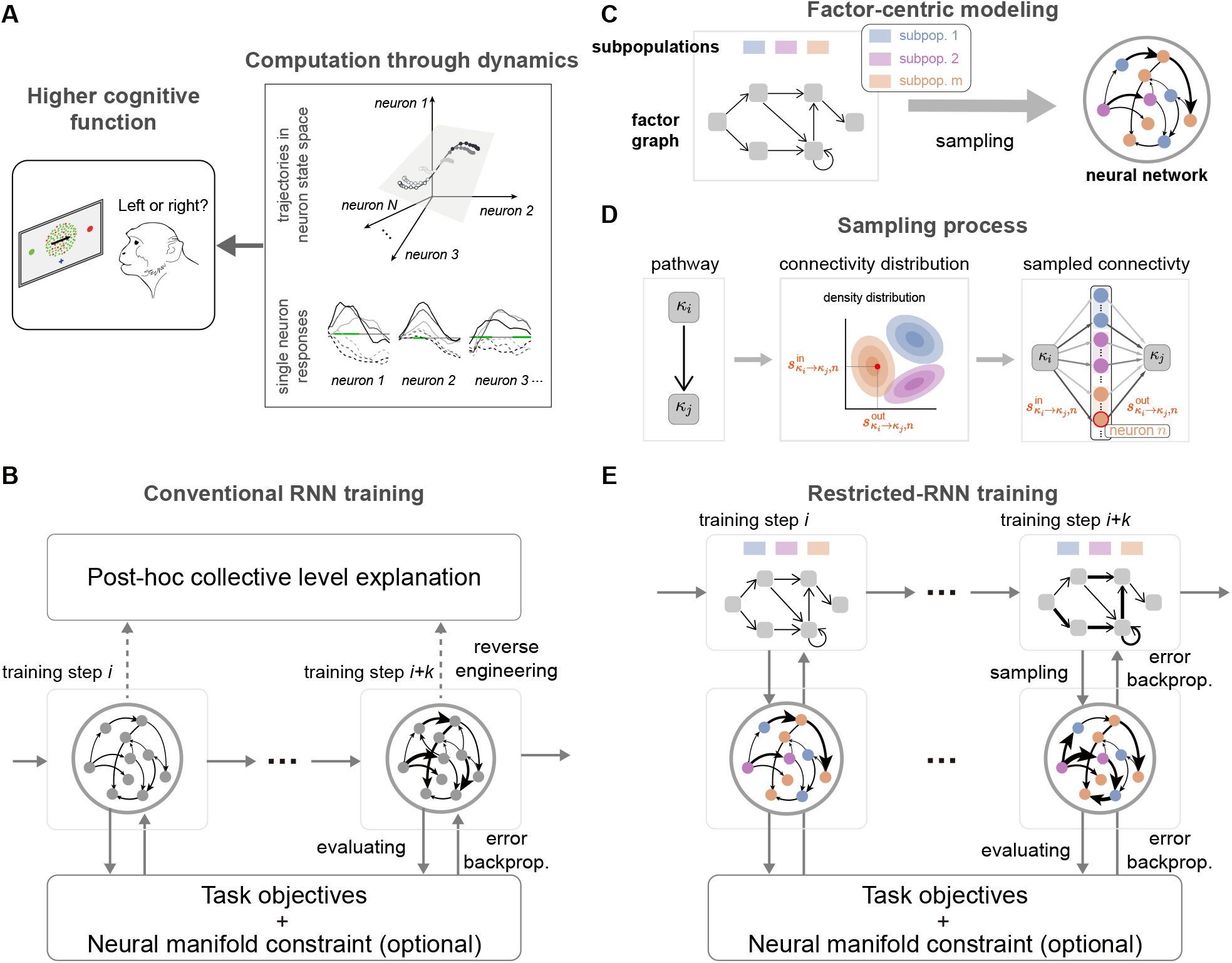
Restricted-RNN—an interpretable factor-centric RNN modeling approach. (A) The computation-through-dynamics (CTD) framework helps reveal circuit mechanisms underlying cognitive functions, accommodating both single-neuron-level activities and collective-level factor representations. (B) The CTD-associated RNN modeling approach (termed as conventional RNN). The conventional RNN posits that the cognitive computations (e.g., factor representations and interactions) emerge from the complex interaction between individual neurons—a neuron-centric view. As such, conventional RNN optimizes the model performance by directly optimizing the connection weights between individual neurons step-by-step through backpropagation. However, the black-box nature of the resulting RNN requires extensive reverse engineering for a post-hoc collective-level interpretation. (C) A factor-centric RNN modeling approach—Restricted-RNN. Restricted-RNN rests on a factor-centric view that centers on directly modeling the interaction between factors. In the form of a collective-level factor graph, Restricted-RNN specifies how the interaction between factors is mediated by multiple neural subpopulations (see panel D for more details). When individual neurons are sampled from neural subpopulations, such a factor graph is then instantiated into a low-rank neural network. (D) Factor communication through neural subpopulations. To send information from one factor (say *k*_*i*_) to another factor (say *k*_*j*_), *k*_*i*_ needs to first go through neurons via input connectivity, each of which then send information to *k*_*j*_ through output connectivity. Importantly, these connectivity weights are not totally random in Restricted-RNN. Instead, they are sampled from a distribution with clusters, each of which corresponds to one distinct neural subpopulation. The input connectivity from *k*_*i*_ to a subpopulation and the output connectivity from that subpopulation to *k*_*j*_ consist of a subpopulation-mediated pathway for the communication from *k*_*i*_ to *k*_*j*_. Note that despite the input and output connectivity weights along each pathway is fixed, how much information can be effectively transmitted from *k*_*i*_ to *k*_*j*_ also depends on the gain of neurons along the pathway. Therefore, the effective coupling strength from *k*_*i*_ to *k*_*j*_ can be potentially modulated by all factors in the graph capable of changing the neural gain (see Methods for more details). (E) Training in Restricted-RNN. Given task objectives and optional neural manifold constraint, instead of optimizing connectivity weights between individual neurons in conventional RNN, we optimize the statistical parameters of connectivity weight distribution in Restricted-RNN through backpropagation by using the reparameterization trick (Kingma & Welling, 2022). In contrast to “black-box” models trained using conventional RNN, endowed by the dynamic mean-field theory, the trained model in Restricted-RNN is fully interpretable (see Methods for more details).

We interrogated the origin of this “black-box” issue in RNNs from a dual-perspective framework—the neuron-centric versus factor-centric views—in cognition (Barack & Krakauer, 2021; DePasquale et al., 2023). Fundamentally, the training of RNNs aims to instantiate task-relevant cognitive factors and simulate their interactions. This objective naturally entails that, within the dual-perspective framework, the interpretation of the model is intrinsically factor-centric. However, the standard training of an RNN optimizes the weights of connections between individual neurons, a process that inherently reflects the neuron-centric view. This mismatch between the neuron-centric view of model training and the factor-centric view of model understanding underlies the lack of model interpretability and related issues in the standard RNN training approach.

Here, we propose a factor-centric RNN modeling approach, termed as Restricted-RNN, capable of removing this mismatch and thereby generating interpretable circuit hypotheses. Restricted-RNN is based upon a factor-centric view of neural computation which regards neurons as the substrate mediating the communication between factors (DePasquale et al., 2023). By introducing the concept “factor communication through subpopulations” (Hirokawa et al., 2019; Hocker et al., 2021; Dubreuil et al., 2022), the dual of “neuron communication through subspaces” (Kohn et al., 2020), we can directly train connectivity weights that mediate the communication between factors (Fig. 1E), which is in sharp contrast to train connectivity weights between individual neurons in conventional RNN (Fig. 1B; Sussillo & Barak, 2013; Yang & Wang, 2020). By doing so, the trained models through Restricted-RNN are automatically endowed with high interpretability as both connectivity training and model understanding are factor-centric (Mastrogiuseppe & Ostojic, 2018; Dubreuil et al., 2022). More importantly, the interpretable nature of Restricted-RNN enables the development of a unified theory to address the core aspects of cognitive control through a common low-dimensional *neural control state space*—offering a compelling geometric view of the ubiquitous control in cognition. We demonstrated the validity of Restricted-RNN through the identification of *de novo* circuit mechanisms underlying both the sequence working memory control (Xie et al., 2022; Chen et al., 2024) and the counter-intuitive firing rate reversal in perceptual decision-making (Okazawa et al., 2021). In both cases, key predictions from the model were validated using monkey neurophysiological data. Together, these results indicate that Restricted-RNN holds the promise to uncover novel circuit mechanisms underlying challenging higher cognition problems.

## Results

### Overview of Restricted-RNN

In contrast to *model the interaction between neurons* in conventional RNN, starting from a factor-centric view of neural computations, Restricted-RNN directly *models the interaction between factors* with a factor graph (Fig. 1C, bottom left). In the graph, each edge indicates a communication pathway between the sender and receiver factors, e.g., *k*_*i*_ and *k*_*j*_, and is formally referred to as the *k*_*i*_ → *k*_*j*_ pathway (Fig. 1D, left). Importantly, this kind of communication pathways between factors is assumed to be mediated by neural subpopulations (Dubreuil et al., 2022; Fig. 1C, top left). That is, to send information from *k*_*i*_ to *k*_*j*_, *k*_*i*_ must first go through individual neurons via input connectivity, which then send the integrated information to *k*_*j*_ through output connectivity (Fig. 1D, right). Moreover, these connectivity weights are not randomly distributed. Instead, they are sampled from a distribution with multiple clusters (Fig. 1D, middle; see Methods for more details), each corresponding to one neural subpopulation, and corroboratively contribute to the information communication along the *k*_*i*_ → *k*_*j*_ pathway. When all pathways in the graph are expanded through the same sampling process, the factor graph then will be instantiated into a low-rank neural network (Fig. 1C, right; see Methods for more details). This pathway-based model construction endows Restricted-RNN with both *great model expressivity* and *high model interpretability*.

The model *expressivity* rests upon two complementary ways in which one factor acts on another— driver and modulator (Dubreuil et al., 2022). A given factor (say *k*_*i*_) can exert its impact on another factor (say *k*_*j*_) as a driver through the *k*_*i*_ → *k*_*j*_ pathway. Alternatively, *k*_*i*_ can also exert its impact on *k*_*j*_ as a modulator through modulating pathways towards *k*_*j*_. How can this modulatory effect occur? In the first glance, given a pathway towards *k*_*j*_ (say *k*_*l*_ → *k*_*j*_), since the input and output connectivity weights along this pathway are fixed, there seems impossible to have a factor away from *k*_*l*_ → *k*_*j*_ such as *k*_*i*_ influence the information transmission along *k*_*l*_ → *k*_*j*_. However, as the information transmission along *k*_*l*_ → *k*_*j*_ is mediated by neurons (Fig. 1D, right), how much information can be effectively transmitted depends on the gain of neurons along the pathway. Therefore, the effective coupling strength of *k*_*l*_ → *k*_*j*_ can be modulated by all factors (including *k*_*i*_) in the graph capable of changing the neural gain (see Methods for more details). Together, these two complimentary roles—driver and modulator—endowed Restricted-RNN with the great fitting power to account for the complex interactions among different factors. Formally, it can be proved that with enough neural subpopulations, Restricted-RNN can approximate any given low-dimensional factor dynamics (see Supplementary Note 4).

The model interpretability rests upon the property that the collective-level connectivity weights (i.e., the statistical parameters of connectivity weights), rather than single-neuron-level connectivity weights (i.e., connectivity weights between individual neurons), can be directly trained in Restricted-RNN (Fig. 1E, see Methods for details). While training single-neuron-level connectivity weights in conventional RNN leads to “black-box” models without a clear collective-level description of factor dynamics (Fig. 1B), training the collective-level connectivity weights in Restricted-RNN naturally leads to models with a highly interpretable collective-level description of factor dynamics (Dubreuil et al., 2022, see Methods for more details). Critically, such a collective-level description encompasses a novel theory for cognitive control, which will be introduced in Fig. 5.

Based upon Restricted-RNN, we then introduce a working pipeline capable of generating hypothesis for a given cognitive problem. The pipeline works in a proposing-and-testing manner: In the proposing step (Fig. 2A, left), a Restricted-RNN with specified graph structure and subpopulation number is proposed; In the testing step (Fig. 2A, right), the collective-level connectivity weights of the proposed Restricted-RNN are trained through backpropagation to test if the proposed circuit hypothesis can solve the given cognitive problem (i.e., fulfill task objectives and neural manifold constraints); If the testing step fails, the proposed hypothesis will be updated through changing either the graph structure (Fig. 2B) or the subpopulation number (Fig. 2C).

**Figure 2.**
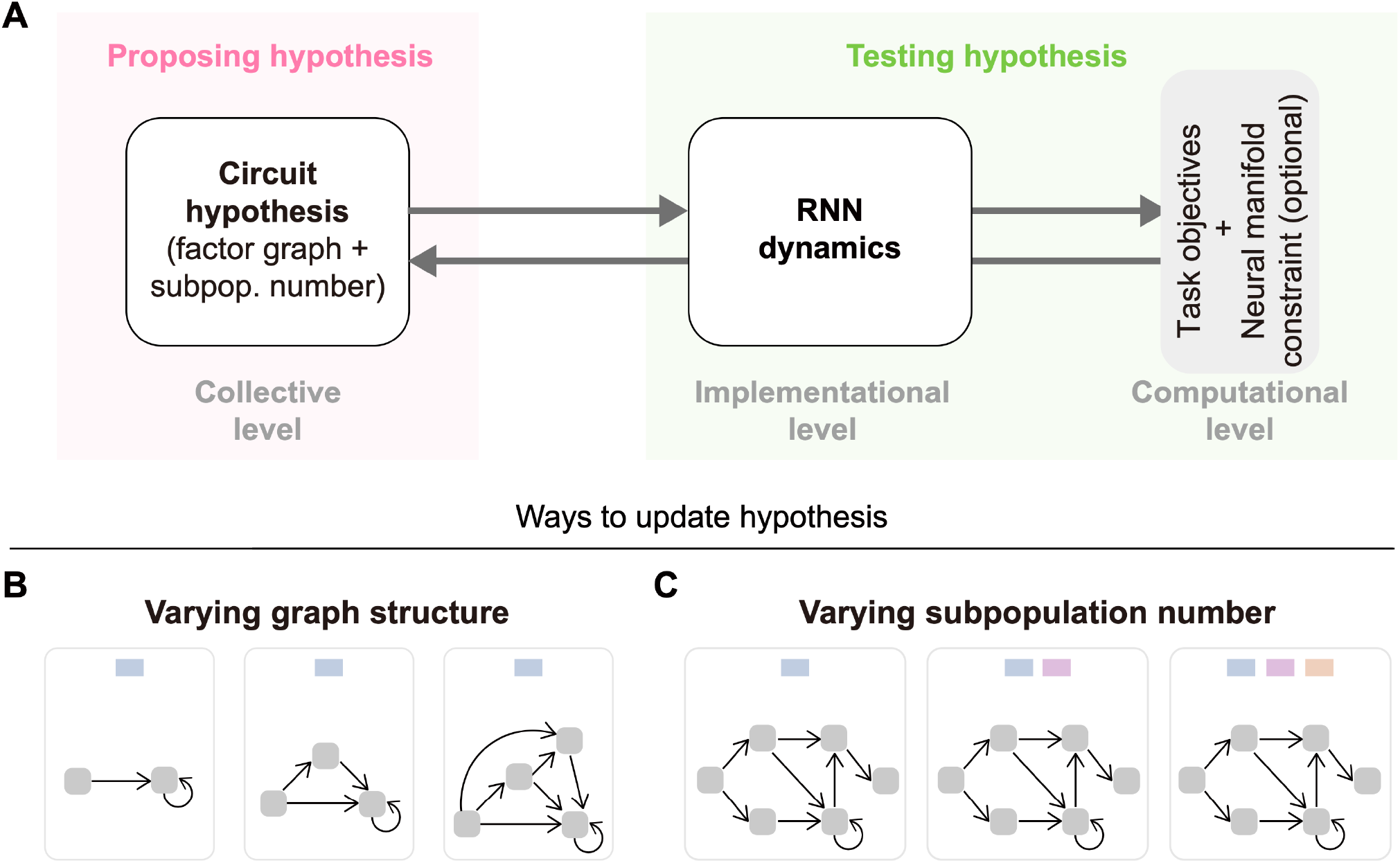
A Restricted-RNN-based working pipeline for hypothesis generation. (A) Given task objectives and optional neural manifold constraint, a Restricted-RNN-based working pipeline is developed to generate the underlying circuit hypothesis. This pipeline works in a proposing-and-testing manner: In the proposing step, a Restricted-RNN with specified graph structure and subpopulation number is proposed; In the testing step, the collective-level connectivity weights of the proposed Restricted-RNN are trained through backpropagation to test if the proposed circuit hypothesis can fulfill task objectives and neural manifold constraint; If the testing step fails, the proposed hypothesis will be updated through changing either the graph structure (as shown in panel B) or the subpopulation number (as shown in panel C). (B) Updating the graph structure in a gradual manner while keeping the subpopulation number fixed. (C) Increasing the subpopulation number in a gradual manner while keeping the graph structure fixed.

In the following, we will validate this pipeline as a powerful circuit hypothesis generator through two representative examples. In the first example, through varying graph structure while keeping subpopulation number fixed, we showed that this pipeline can generate novel circuit hypothesis for the counter-intuitive firing rate reversal in perceptual decision-making (Okazawa et al., 2021), with the key predictions being confirmed by monkey parietal cortex data (Fig. 3). In the second example, through varying subpopulation number while keeping graph structure fixed, we demonstrated that this pipeline can generate *de novo* circuit hypothesis for the sequence working memory control problem (Xie et al., 2022; Chen et al., 2024), with the key predictions being experimentally confirmed by monkey frontal cortex data (Fig. 4).

**Figure 3.**
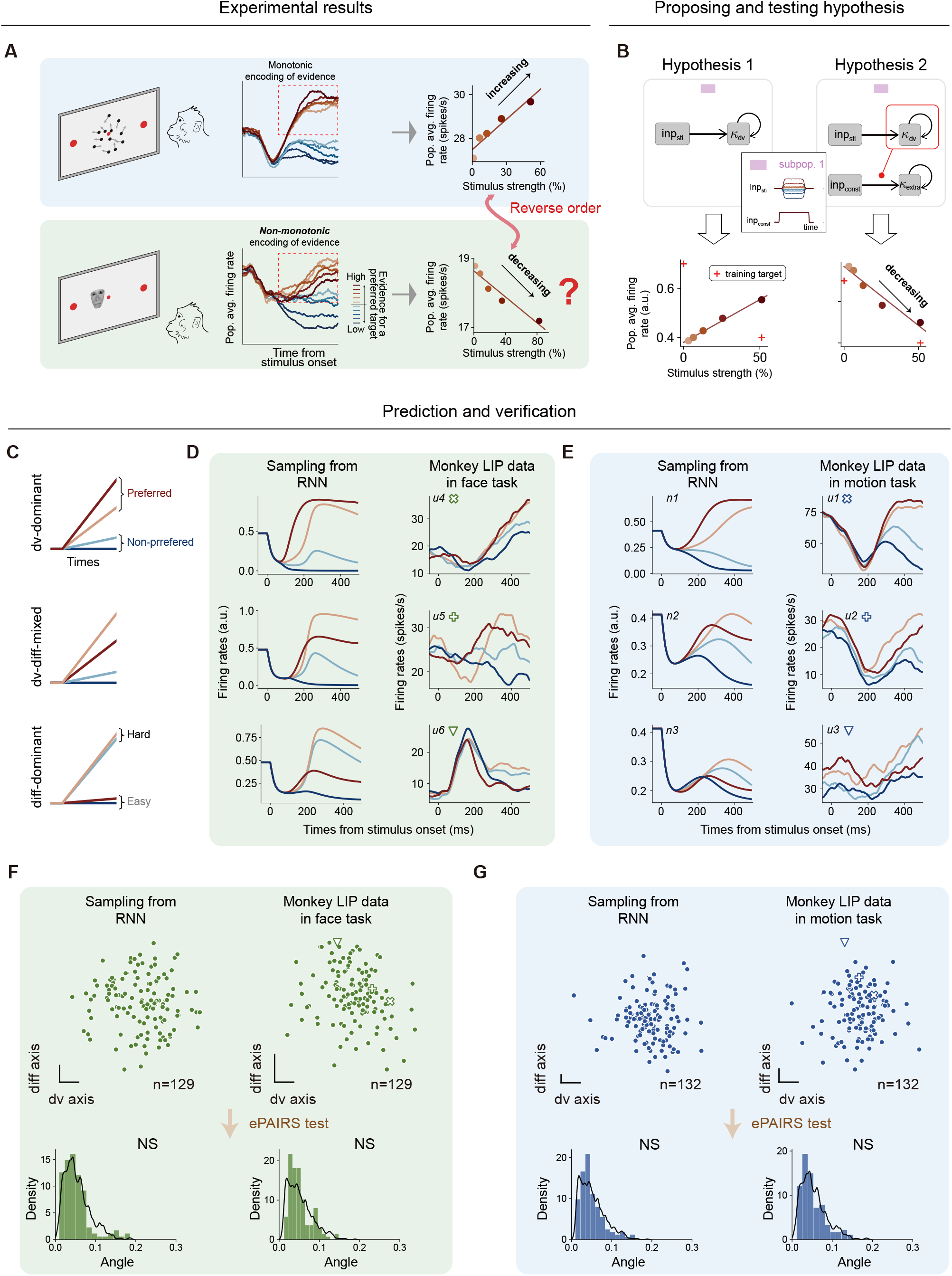
Restricted-RNN unraveled a novel circuit mechanism underlying mean firing rate reversal in perceptual decisions. (A) Two perceptual decision-making (PDM) tasks (Okazawa et al., 2021): (upper) the motion PDM task, where monkey responds based on the overall movement of random dots. (bottom) The face PDM task, where monkey responds according to how closely a presented face resembles either a human or a monkey face. Previous research found that, unlike the motion PDM task, neurons in the monkey lateral intraparietal area (LIP) in the face PDM task display an counter-intuitive firing rate pattern—mean firing rate reversal—as shown by non-monotonic tuning with respect to the evidence for the neurons’ preferred target (middle) and decreasing mean firing rate with respect to increasing input stimulus strength (right). (B) Generating hypothesis for meaning firing rate reversal through varying the factor graph while keeping the subpopulation number fixed. Starting from the basic circuit (Hypothesis 1) that only involves one factor *k*_dv_ for the evidence accumulation, we systematically tested several hypotheses (see Methods) and found that it is necessary to include an additional factor *k*_extra_ (Hypothesis 2). The final collective circuit has two factors: *k*_dv_ integrates the input evidence, while *k*_extra_ receives and accumulates a constant input, exhibiting a stimulus difficulty-like code due to the modulation effect of *k*_dv_ on the *inp*_const_ → *k*_extra_ pathway (red line with circle). (C-G) Model predictions and verification. Restricted-RNN model predicts that both the motion and face PDM tasks consist of three types of neurons: i) dv-dominant neurons that monotonically respond to evidence strength, ii) diff-dominant neurons that respond to the absolute evidence (difficulty) of stimuli, and iii) dv-diff-mixed neurons that display mixed tuning and non-monotonical responses to evidence strength (C). We found neurons corresponding to all the three types from the face PDM task (D) and motion PDM task (E). In each panel, the left column shows Restricted-RNN neurons, and the right column shows neurons from monkey LIP data. From top to bottom, the example neurons belong to the dv-dominant, dv-diff-mixed, and diff-dominant type, respectively. Panels (F) and (G) show the population structure in the face and motion PDM tasks, respectively. Population structure is characterized by projecting neurons onto the *k*_dv_ and *k*_extra_(or *k*_diff_) axes (upper plot in each panel), reflecting neurons’ roles in coding these factors. In these plots, each point stands for a neuron from either Restricted-RNN (left) or Monkey LIP data (right). In all cases, there exist only one neural subpopulation (ePAIRS test, see Methods).

**Figure 4.**
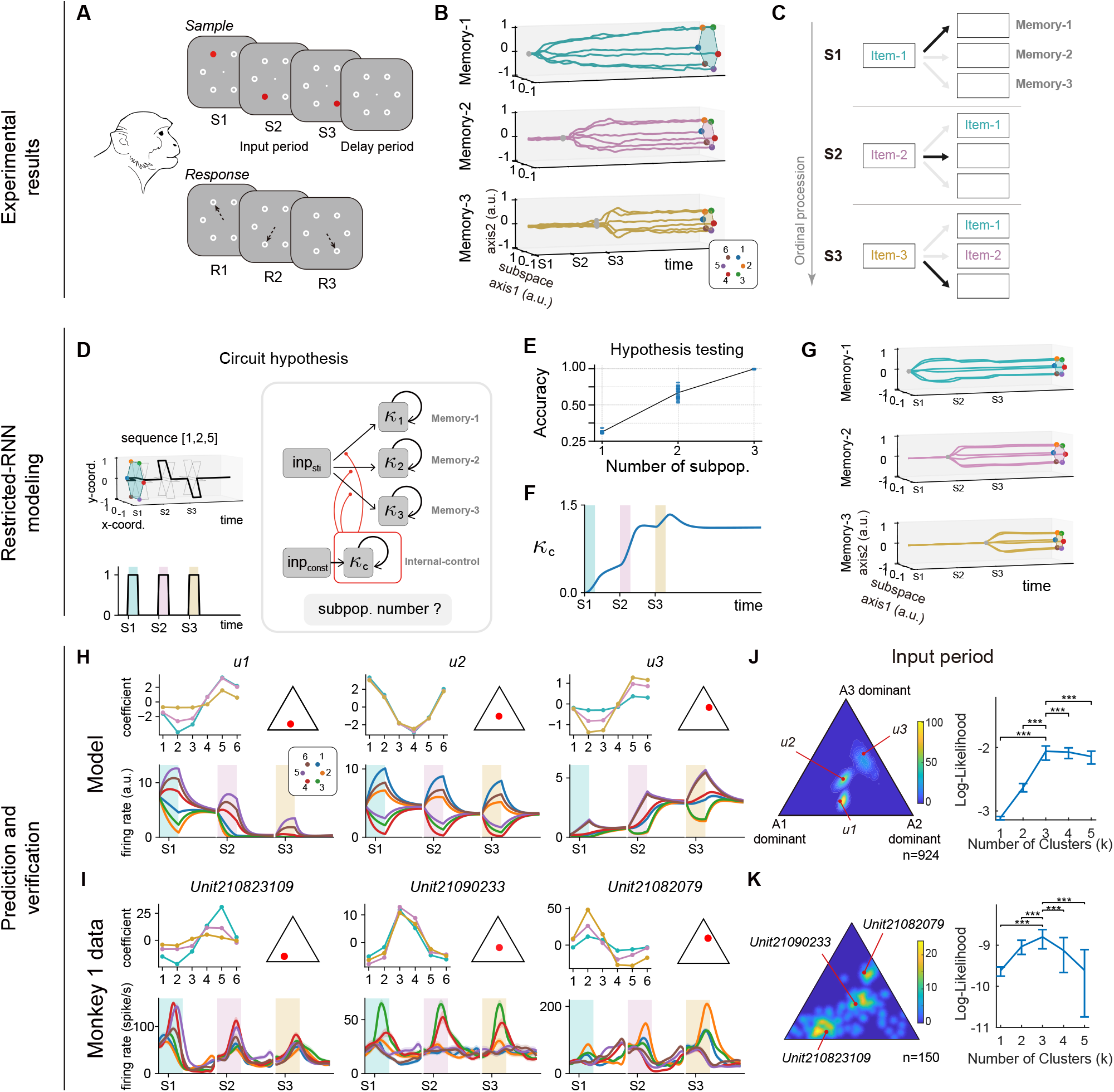
Restricted-RNN modeling for sequence working memory task predicts the minimal subpopulation structure underlying memory gating. (A) Sequence working memory (SWM) task (Chen et al., 2024). The monkey receives a sequence of stimuli (of length 1-3) each being a location drawn from a hexagon. The task requires the monkey to remember both the locations and their order for a variable delay period (Delay) and then reports the locations in the presented order. (B) Neural trajectories in the memory subspaces for the length-3 SWM case. Trajectories from the input period to the delay period are shown, colored according to the dominant ordinal rank (rank 1, 2, 3 for memory subspace 1, 2, 3, respectively). (C) A summary of the memory gating process of SWM. At first the gateway to memory subspace 1 opens while the other two are closed, so that the first item in the sequence enters memory subspace 1. Then this gateway shuts down and the one to memory subspace 2 opens. Therefore, the second item enters memory subspace 2. Finally, the same process guides the last item entering memory subspace 3. (D-G) Generating hypothesis for SWM gating through varying subpopulation number while keeping the graph structure fixed. (D) Factor graph based on previous findings in (C). There are three factors (*k*_1_, *k*_2_, *k*_3_) for integrating the memory singals of the three ranks, respectively. They share the same stimulus input. In addition, the factor *k*_*c*_ receives an input indicating the presence of input stimuli so that *k*_*c*_ can act as an internal ordinal rank signal to modulate all the pathways connecting the sensory input with sequence working memory (red line with circle). (E) We tested hypotheses with increasing number of subpopulations and found that it requires at least three subpopulations for Restricted-RNN to accomplish the length-3 SWM task. (F) Factor *k*_*c*_ exhibited a temporal profile encoding the ordinal rank of the sequence. (G) Neural trajectories in the memory subspaces of the trained Restricted-RNN, which replicates the experimental observation in (B). (H-K) Model predictions and verification. (H) and (I) show example neurons of Restricted-RNN and Monkey PFC data. During the input period, item-selective Restricted-RNN neurons exhibit stable tuning phases across different ordinal ranks (H). Similar patterns can be found in Monkey PFC data (I). Therefore, the amplitude distribution of different ordinal ranks (distribution of *A*_1_, *A*_2_, *A*_3_) captures the essence of neuron tuning pattern during the input period. We visualize (see Methods) the amplitude distribution across neurons in panels (J) and (K), using color to indicate neuron density—brighter colors represent higher densities. Clustering analysis (see Methods) reveals that there are three subpopulations in Restricted-RNN model (J), and the same is true for Monkey PFC data (K). One-side Wilcoxon signed-rank test with Benjamini-Hochberg FDR correction for multiple comparisons ***P<0.001.

### Restricted-RNN uncovered a *de novo* neural mechanism explaining the counterintuitive firing rate reversal in macaque parietal cortex

One of the best ways to validate a new approach is to test if it can explain counterintuitive phenomena. We therefore asked whether Restricted-RNN could account for a recent finding of counterintuitive, task-dependent neural geometry in the monkey parietal cortex during perceptual decision-making tasks (Okazawa et al., 2021). In this study, the mean firing rates of lateral intraparietal (LIP) neurons were found to encode sensory evidence non-monotonically (i.e., mean firing rates were reversed with respect to the evidence for the neurons’ preferred target) during a face discrimination task (Okazawa et al., 2018, 2021; Fig. 3A, bottom panel). This contradicts classical empirical findings of monotonic evidence encoding in LIP during motion discrimination tasks (Shadlen & Newsome, 2001; Gold & Shadlen, 2007; Churchland et al., 2008; Bennur & Gold, 2011; Purcell & Kiani, 2016; Fig. 3A, top panel), as well as classical bistable attractor models of perceptual decisions (Wang, 2002; Mazurek et al., 2003; Lo & Wang, 2006; Wong & Wang, 2006; Purcell et al., 2010; Wimmer et al., 2015). The study showed that these contradicting results arise because sensory evidence is encoded on a task-dependent, curved manifold in neural state space (Okazawa et al., 2021), in which both decision variable (dv) and stimulus difficulty (diff, the unsigned strength of sensory inputs) were encoded (Fig. S3-1). However, the key question of how this curved manifold and the reversal of firing rates arise from a biological neural circuit remains unanswered, providing an ideal case for testing the validity of Restricted-RNN.

In the following, we demonstrated that Restricted-RNN offers a systematic solution to addressing this challenging issue. Basically, this issue can be translated into the following computational problem: what is the minimal circuit model that can simultaneously perform the perceptual decision-making and satisfy the firing rate reversal manifold constraint? To start with, we first proposed the simplest perceptual decision-making circuit (Fig. 3B, middle panel; Fig. S3-2, A and D; termed as H1; Dubreuil et al., 2022), with only one pathway sending the input variable *inp*_sti_ to the decision variable *k*_dv_. In the testing step, after model training, we found that H1 can generate a curved manifold (Fig. S3-3A) but cannot reproduce the reversal of mean firing rates (Fig. 3B, left panel). What is the underlying mechanism? The high interpretability of Restricted-RNN enabled us to answer this question. First, while the latent variable *k*_dv_ in H1 indeed forms a straight line, the observed neural activity is not the latent variable *k*_dv_ per se. Instead, it corresponds to a nonlinear transformation (i.e., the nonlinear activation function) of *k*_dv_, effectively transforming the straight line into a curved manifold (see Methods for more details; Okazawa et al., 2021). Second, while this simple nonlinear transformation can produce a curved manifold, it cannot yield the reversal of mean firing rates. This is because as a monotonic-increasing function, the nonlinear activation function alone cannot change the monotonicity of mean firing rate (see Methods for more derivations).

To replicate this non-trivial firing rate reversal, we then increased the model complexity by adding an additional latent factor (*k*_extra_ ) with extra pathways while keeping the number of subpopulations equal to one. However, we found that none of these models could reproduce the firing rate reversal (Fig. S3-3B, left and middle panels). Inspired by the classical Wong-Wang model (Wong & Wang, 2006), we then introduced a constant input *inp*_const_, independent of the stimulus strength, to mimic the overall excitation of neurons during the task, but found that this kind of circuit still failed to generate the firing rate reversal (Fig. S3-3A, right panel). Surprisingly, when the pathway *inp*_const_ → *k*_extra_ was added to the model structure (termed as H2), the firing rate reversal could be conveniently reproduced with only one subpopulation (Fig. 3B, right panel; see Methods for more details) and *k*_extra_ in the trained model behaved like a difficulty variable— encoding the absolute value of input strength (Figs. S3-2, B and E; see Fig. 5B for a novel geometric interpretation for how this model works). This H2 also led to the following novel testable predictions at both single-neuron and collective levels.

First, H2 predicts that the decision and difficulty variables are represented within a single neural subpopulation. Depending on their relative encoding strengths, individual neurons should exhibit three patterns: a dv-dominant (monotonic) pattern (Figs. 3, C and D, top panel), a diff-dominant pattern (high firing for hard trials, Figs. 3, C and D, bottom panel), and a dv-diff-mixed (non-monotonic) pattern (Figs. 3, C and D, middle panel). Through re-examining the parietal cortex neuron data (Okazawa et al., 2021), we indeed found all three single neuron firing patterns in face task (Fig. 3D, right panel; Fig. S3-4, A and C), verifying our model predictions.

Second, at the collective level, our modeling result predicted that one neural subpopulation suffices to explain the experimentally observed neural geometry. To test this prediction, we performed the ePAIRS analysis (Raposo et al., 2014; Hirokawa et al., 2019; Dubreuil et al., 2022) and found that these seemingly diverse neural firing patterns in the face task could be well-explained only by one neural subpopulation (Fig. 3F; Fig. S3-6, A; see Methods for details), perfectly aligning with H2 and thereby verifying our collective-level model prediction. The same conclusion held with a likelihood-based clustering method (Fig. S3-5A and Fig. S3-6C).

Importantly, when the same single-neuron and population analysis were applied to the motion task, we found the almost same firing patterns: there were three different kinds of single-neuron firing profiles (Fig. 3E, right panel and Fig. S3-4, B and D) and one functional cluster (Fig. 3G, right panel; Fig. S3-5, B; Fig. S3-6, B and D). We shall emphasize that while the dv-diff-mixed pattern was reported in the face task (Okazawa et al., 2021), the existence of both dv-diff-mixed and diff-dominant single neurons in LIP in the motion task is particularly surprising as LIP neurons in this task has been traditionally considered to be of dv-dominant type (Gold & Shadlen, 2007). From the modeling perspective, the existence of diff-dominant single neuron cannot be explained by H1, thereby challenging the canonical circuit model for the motion task (see Methods for more details). Interestingly, we found that the same H2 but with a set of different collective connectivity parameters can conveniently explain all these patterns (Fig. 3E, left panel; Fig. 3G, left panel; Fig. S3-5, B; Fig. S3-6, B and D). Therefore, H2 provided a unified yet simple neural mechanism for explaining the neurophysiological data in both motion and face decision-making tasks.

Taken together, our systematic exploration with Restricted-RNN not only provided a minimum, interpretable circuit hypothesis underlying the counterintuitive firing rate reversal in monkey LIP but also revealed novel properties of neural responses even in the classical motion task, showcasing the strength of Restricted-RNN in exploring circuit mechanisms from experimental data in cognition tasks.

### Restricted-RNN uncovered a *de novo* neural mechanism underlying sequence working memory control

To further test the validity of Restricted-RNN, we then applied it to a recent sequence working memory experiment (Chen et al., 2024; Xie et al., 2022). In this experiment, monkeys were presented with a spatial sequence and required to reproduce the sequence after a variable delay period (Fig. 4A). Neural data analysis revealed that spatial items in different ordinal ranks were routed to the corresponding rank subspaces in monkey frontal cortex upon the stimulus presentation and stably maintained throughout the delay period (Fig. 4B). The remaining open question is how such a precise information control—opening the pathway from the input to the right rank subspaces while closing other input pathways at each input period (Fig. 4C)—is achieved in monkey frontal cortex. There are a few proposals in the literature (Botvinick & Plaut, 2006; Botvinick & Watanabe, 2007). For example, one recent work proposed using memory rotation to solve the SWM control problem (Fig. S4-1A; Whittington et al., 2024). However, further neural data analysis showed that none of these proposals are well-supported by the frontal neural data (Fig. S4-1B). We then asked if Restricted-RNN can generate novel circuit mechanisms consistent with the frontal neural data.

Enlighted by the experimental findings of length-3 sequence working memory representations (Chen et al., 2024), we introduced four factors, including one spatial location input factor (*inp*_*sti*_) and three working memory factors (*k*_*r*_, *r* = 1,2,3), into the circuit hypothesis (Fig. 4D). There are three feedforward pathways (*inp*_sti_ → *k*_*r*_, *r* = 1,2,3) sending the spatial location input information to working memory factors and three recurrent pathways (*k*_*r*_ → *k*_*r*_, *r* = 1,2,3 ) for memory maintenance (Fig. S4-2, see Methods for more details). In addition, to monitor the task stage procession (i.e., rank-1, -2 or -3 period), we introduced an integrator variable *k*_*c*_ that can integrate a spatial-location-independent input *inp*_const_ across the sample period through the *inp*_const_ → *k*_*c*_ pathway. This task-stage-monitoring signal *k*_*c*_ is hypothesized to modulate the information pathway from spatial location input factor to different working memory factors (i.e., *inp*_sti_ → *k*_*r*_, *r* = 1,2,3) through subpopulation-based gain modulation (Fig. 4D, red line with circle). With this factor graph structure, we then ask the following question: what is the minimal number of subpopulations required to perform this SWM task? In the testing step, by gradually increasing the number of subpopulations starting from *M* = 1, we identified that the minimal number of subpopulations is 3 for the length-3 sequence working memory control problem (Fig. 4E). In this minimal model, we verified that factor *k*_*c*_ indeed acted as a task-stage-monitoring signal (Fig. 4F) while the three working memory factors exhibited the similar dynamics with the experiment (Fig. 4G).

Then, what did these three subpopulations look like? An intuitive possibility (Figs. S4-3, A and C) is that each subpopulation is activated only in one input period (e.g., subpopulation 1 for rank-1 period). Simply examining the neural firing pattern in different input periods, however, rejected this possibility (Fig. 4I). Instead, most of neurons were activated during multiple input periods. To better characterize this neural firing pattern, for a given neuron, we introduced *A*_*r*_ and *θ*_*r*_ (*r* = 1,2,3), of which *A*_*r*_ and *ϕ*_*r*_ quantified the activation amplitude and the preferred location with respect to the input during the *r*-th input period, respectively (Fig. S4-3B; see Methods for more details). Through examining the preferred location during the input periods, we found that most of neurons in the model showed the same location preference at different input periods—a gain modulation profile (Fig. 4H and Fig. S4-4, A and C). As a control, we also examined the preferred location of these neurons during the delay period and found that most of neurons showed different location preferences during this memory period (Fig. S4-4, B and D). To further investigate the collective-level pattern of neural firing, we collected all activation amplitudes at different input periods, i.e., (*A*_1_, *A*_2_, *A*_3_), and examined the neuron density distribution in the space of (*A*_1_, *A*_2_, *A*_3_). We found an apparent clustering structure is exhibited in this space (Fig. 4J). As expected, this distribution is best explained by three clusters using likelihood method (see Methods for more details; regarding how this model works, see Fig. 5 for a novel geometric understanding).

**Figure 5.**
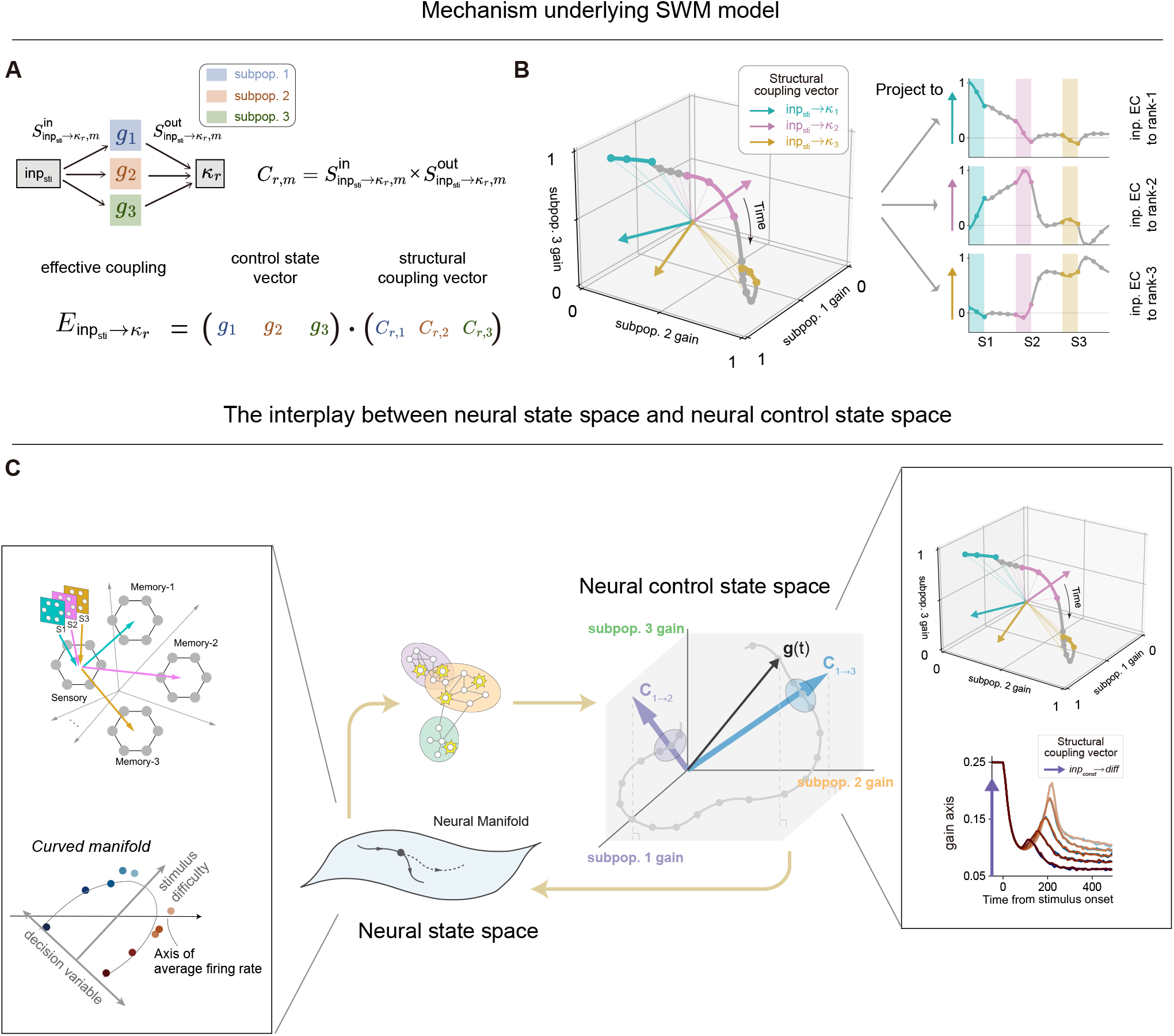
The control state space provides a unified framework for interpreting the control process. (A-B) The control state space description and mechanism underlying memory gating in SWM model. (A) At collective-level, the effective coupling strength *E*_*p*_ along a given pathway *p* (say *inp*_*sti*_ → *k*_*r*_) in SWM model equals to the inner product of two quantities: the structural coupling vector ***C***_*p*_ = (*C*_*p*,1_, ⋯, *C*_*p,M*_) and the subpopulation gain vector ***g*** = (*g*_1_, ⋯, *g*_*M*_). Here, each structural coupling component *C*_*p,m*_ equals to the multiplication of the collective-level output input connectivity strength 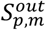 and input connectivity strength 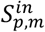, while each gain component *g*_*m*_ equals to the average gain of neurons in subpopulation *m* (see Methods for more details). This inner product formulation endows the model with a geometric interpretation for the underlying control mechanism: how much information can be transmitted along a given pathway *p* is determined by the degree of alignment between the structural coupling vector ***C***_*p*_ and the subpopulation gain vector ***g*** (termed control state vector) in a new state space (namely the control state space) with each axis representing one subpopulation gain. (B) The evolution of control state vector (grey line) in the newly introduced control state space. During input periods S1, S2 and S3, the control state vector exclusively aligns well with rank-1, rank-2 and rank-3 structural coupling vectors, respectively, thus opening the gateway to corresponding memory subspaces successively. (C) The control state space provides a concise and intuitive way to understand how the flow-field in the neural state space is formed and guide the evolution of neural representations. Specifically, neural connectivity establishes structural coupling vectors, and the arrangement of the control state vector relative to these coupling vectors in the control state space governs the information flow (right box). This interaction influences the evolution of representations in the neural state space (left box), altering the gain of individual neurons and thereby affecting the control state vector. This description also can be applied to elucidate the mechanism underlying the emergence of difficulty representation in the PDM model. Despite there is no direct difficulty input, the control state scalar evolves differently with different absolute stimulus strengths (right box), thereby influencing how factor *k*_extra_ integrates the stimulus-irrelevant input along the *inp*_const_ → *k*_extra_ pathway. The difficulty representation exhibited by *k*_extra_ (left box) inherits from the symmetric structure of control state scalar in the control state space (right box, see Methods for more details).

We then asked if there is any experimental evidence for supporting such a model. Following the same analysis, we found that most single neurons in monkey frontal cortex indeed showed a gain-modulation activity pattern (Fig. 4I and Figs. S4-4, E and G) during the input period and a preference shift pattern during the delay period (Figs. S4-4, F, H, I and J), confirming the model prediction. Furthermore, the same clustering analysis confirmed the existence of a subpopulation structure with at least three clusters in the space of (*A*_1_, *A*_2_, *A*_3_) in monkey frontal cortex (Fig. 4K, Fig. S4-5B and Figs. S4-6 A and B for monkey 1; see Fig. S4-5, A and C and Figs. S4-6 C and D for monkey 2), which validated the model predictions. Critically, the presence of multiple gain-modulated firing patterns in monkey frontal cortex during the input period indicates a radically different picture regarding the SWM gating process: instead of interpreting the transient responses during the input period with a shared entry subspace (Chen et al., 2024), these transient responses should be interpreted as the footprint of ordinal-rank-dependent gain modulation, a dynamic signature of subpopulation-mediated SWM control mechanism.

Together, this example demonstrated that Restricted-RNN can generate novel circuit hypothesis and make testable predictions. Notably, these minimal model-based predictions were well-supported by monkey frontal data, which is surprising given the complexity of biological data. This result suggests that despite staying in a relatively abstract level in terms of factors, subpopulations and pathways, our modeling framework can still capture key aspects of biological neural computation.

### Restricted-RNN endowed us with a unified theory to understand the hidden control representation

Through these examples, we have demonstrated Restricted-RNN as a powerful hypothesis generator. In fact, as a modeling approach grounded in rigorous theoretical basis, Restricted-RNN can also endow us with a unified theory to explain the disparate phenomena across different tasks. Let us first recall the computation-through-dynamics framework (Vyas et al., 2020; DePasquale et al., 2023), in which hypothesis regarding cognitive computation is expressed in terms of flow-field shaping the trajectory of factors in the high-dimensional neural state space. In general, it is hard to get an intuitive understanding of the flow-field in a high-dimensional space, the key to the “black-box” issue. Interestingly, the interpretable nature of Restricted-RNN can provide us a valuable means for understanding how the flow-field itself is formed to perform the computation required by the task.

In the example of SWM control here, understanding how the flow-field shapes the trajectory of the working memory factor *k*_*r*_ during the input period is equivalent to modeling the effective coupling strength of the pathway *inp*_sti_ → *k*_*r*_. In the mean-field limit of Restricted-RNN, this coupling strength can be concisely expressed as the inner product between the structural coupling vector (*C*_1_, *C*_2_, *C*_3_)^*T*^ of pathway *inp*_sti_ → *k*_*r*_ and the subpopulation gain vector (*g*_1_, *g*_2_, *g*_3_)^*T*^, of which 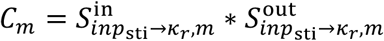 (*m* = 1,2,3) stands for the structural coupling strength mediated by the *m*-th subpopulation while *g*_*m*_ is the average gain of the *m*-th subpopulation (Fig. 5A and Fig. S5-1; see Methods for more details). When introducing a new state space (named as control state space) with each axis standing for the gain of one distinct subpopulation, such an inner product formula then can be interpreted in a geometric manner: 1) for each trial, the trajectory of the state-dependent subpopulation gain vector (named as control state vector) in this control state space represents the dynamic control state of the network; 2) the overlap between this dynamic control state vector and the static structural coupling vector of a given pathway determines the effective coupling of the pathway.

By plotting out both the static structural coupling vectors of three input pathways *inp*_sti_ → *k*_*r*_ (*r* = 1,2,3) and the dynamic control state vector through the whole trial in this new control state space (Fig. 5B, left panel), we found that 1) during the rank-1 stimulus period (cyan trajectory), the dynamic control state vector stays in a subregion with-aligned with the structural coupling vector of the *inp*_sti_ → *k*_1_ pathway; 2) with the precession of task stage, the dynamic control state vector enters in a subregion with-aligned with the structural coupling vector of the *inp*_sti_ → *k*_2_ pathway during the rank-2 stimulus period (purple trajectory); 3) as expected, during the rank-3 stimulus period (yellow trajectory), the dynamic control state vector enters in a subregion with-aligned with the structural coupling vector of the *inp*_sti_ → *k*_3_ pathway. A more quantitative visualization can be obtained by projecting the dynamic control state vector into the three structural coupling vectors, revealing a substantial overlap with the structural coupling vector of the relevant pathway while near-zero overlaps with other irrelevant structural coupling vectors at each input period (Fig. 5B, right panel)—the desired geometric property required by sequence working memory control (Fig. 4C). Notably, this geometric portrait also provides an intuitive understanding of why three subpopulations are required for the length-3 sequence working memory control problem: three dimensions are the minimal requirement to have the dynamic control state vector be overlapping with one structural coupling vector while simultaneously orthogonal to the other two structural coupling vectors.

Importantly, such a set of analyses can be equally applied to the perceptual decision-making model. For the perceptual decision-making model, the major issue is to understand how the difficulty variable emerges from a circuit without explicitly modeling the difficulty variable (Fig. 3B, right panel). To resolve this issue, we applied the same set of analyses to the perceptual decision-making model. In this model, as there is only one subpopulation, the associated new state space is of one dimension. By plotting out the trajectories of the control state (i.e., the subpopulation gain) with different stimulus strengths, we found that upon the onset of stimulus, the subpopulation gains with larger absolute stimulus strengths dropped faster than those with smaller ones, resulting in a difficulty-like code (Fig. S5-2). According to the model, as *k*_diff_ represents the code of *inp*_const_ (a constant code independent of stimulus strengths) multiplied by the effective coupling strength of *inp*_const_ → *k*_diff_ pathway, this explains why *k*_diff_ behaves like a difficulty variable. Together, this set of analyses provides a mechanistic understanding for the emergence of difficulty variable in perceptual decision-making tasks.

Taken together, we propose this new state space, named as *the neural control state space*, as an important complement to the concept of the predominant neural state space (Fig. 5C). In the neural state space, the trajectory of multiple factors can be conveniently investigated, providing a geometric understanding regarding the factor representation but leaving the issue of how these factor representations are formed largely open. Augmented with the subpopulation structure (which is verified in both SWM and decision-making experiments here), the neural control state space provides the much-needed concept to account for the intriguing interactions between factors and thereby explains how the trajectories of multiple factors are formed.

## Discussion

As a deep-learning-based dynamical model, RNN naturally implements the computation-through-neural-dynamics framework and plays a major role in generating circuit mechanism hypothesis in systems neuroscience. However, it also inherits the “black-box” property from deep-learning approach. Through integrating the multi-level descriptions of neural systems and leveraging the recent theoretical progress in neural computation, Restricted-RNN provides an alternative factor-centric modeling approach featuring high model interpretability. The validity of Restricted-RNN in novel circuit hypothesis generation was demonstrated through a variety of macaque cognitive tasks, with the key derived predictions being confirmed by monkey neurophysiological data. Critically, based on Restricted-RNN, a new concept—namely the neural control state space—was proposed to provide a unified geometrical understanding for the ubiquitous control in cognitive processes. Together, these results strongly demonstrated the great promise of Restricted-RNN in generating novel interpretable circuit mechanism hypothesis for challenging higher cognitive problems.

### Post hoc reverse-engineering versus theory-based training

In conventional RNN, each connection weight is trained, endowing the model with great fitting power (with about 10,000 free parameters for an RNN with 100 neurons). While many insights can be gained from these trained models through a variety of reverse-engineering approaches, it is difficult to understand a model with 10,000 free parameters in general. Crucially, this black-box issue could be amplified in challenging problems. For example, recent work showed that conventional RNN failed to reproduce the individual variability of neural computations underlying flexible decisions (Pagan et al., 2025). Instead of training each connection weight between individual neurons, Restricted-RNN directly trained “the collective-level connectivity weights” through introducing a novel generative model for connectivity matrix. By doing so, Restricted-RNN endowed the trained model with high interpretability and conveniently generated data-compatible circuit models for a variety of cognitive tasks.

### Neural state space versus neural control state space

Neural state space has been playing a dominant role in revealing the dynamic evolution of task-related factors (Cunningham & Yu, 2014; Vyas et al., 2020). In other words, it provides the appropriate concept for characterizing the representation of factors, which is undeniably important. However, it is conceivable that the emergence of these factor representations involves a delicate control process that enables a contextual dependent information flow regulation required by the task. What is the right language to describe such a delicate control process remains unknown (Miller & Cohen, 2001; Cohen, 2017; Badre et al., 2021; Panichello & Buschman, 2021; MacDowell et al., 2022; Stine & Jazayeri, 2025). As demonstrated in this work, introducing the neural control state space enabled us to address the following key issues: 1) How is control represented in a neural system? What is the dimensionality of control representation? How does control representation get updated? How does control representation regulate the information flow? Therefore, we proposed that the neural control state space may provide the appropriate concept for accounting for the delicacy of control representation. Further systematic investigation is warranted to better test the generality of this new concept.

### Factor-centric view versus neuron-centric view

Whether brain computation is factor-centric or neuron-centric is hotly debated (Barack & Krakauer, 2021). As Restricted-RNN is task-driven, it naturally adopted the factor-centric view—regarding neural populations as the substrate mediating the interactions among different task-related factors. The introduced pathway-based generative model for connectivity matrix is a natural way to account for the interactions among different factors (Fig. 1C). In fact, in Restricted-RNN, the two views are not contradictory but complimentary (Fig. S5-3): factor interact with each other through neural populations while neural populations communicate with each other through subspace (Kohn et al., 2020; Semedo et al., 2019).

### Restricted-RNN as a systematic hypothesis generator

To be qualified as an ideal hypothesis generator, an approach should be able to systematically explore the hypothesis space. Previous works have showed that conventional RNN may not be able to explore the whole hypothesis space (Pagan et al., 2025). The pathway-based generative model of connectivity introduced here provides a systematic approach to identify the minimal circuit model in terms of the number of connectivity ranks, subpopulations and pathways. By identifying *de novo* circuit mechanisms for both sequence working memory and counter-intuitive firing rate reversal, we demonstrate the validity of this systematic approach, strongly supporting Restricted-RNN as systematic hypothesis generator.

### Connection with classical biophysical models

Classical biophysical models (Compte et al., 2000; Machens et al., 2005; Wong & Wang, 2006), such as Wong-Wang model mentioned here, provide enormous insights into the neural mechanisms underlying various cognitive processes. The model generated from Restricted-RNN provides an alternative description for these cognitive processes at an abstract level in terms of connectivity ranks, subpopulations and pathways. Despite this abstraction, this kind of description still preserves the key ingredients closely linking with neural physiological data, as demonstrated in all of examples presented here. The exhibited parsimony of such an abstract description may provide insights into more complex cognitive process such as sequence working memory manipulation (Tian et al., 2024).

In his famous essay “More is Different”, Philip Anderson argued that “at each level of complexity entirely new laws, concepts, and generalization are necessary” (Anderson, 1972). Consistent with this general concept, the three-level analysis is proposed by David Marr and colleagues to account for the multi-level nature of complex neural systems (Marr, 1982). However, to date, the firm connections between these three levels are yet to be established. Here, by introducing a novel mathematical language in terms of connectivity ranks, subpopulations and pathways, we established such a much-needed link between these three levels and provided a systematic approach towards generating novel circuit hypotheses for challenging higher cognition problems with high dimensionality and high complexity in a learnable and interpretable fashion.

## Acknowledgement

We thank Drs. Srdjan Ostojic, Xuexin Wei, Albert Compete and Cheng Xue for their critical comments on the manuscript.

## Methods

### The Restricted-RNN modeling framework

In this section, we will elaborate on the details of the Restricted-RNN framework, including 1) *a novel pathway-based generative model*, 2) *the pathway-based factor graph and associated RNN*, and 3) *the factor dynamics in the mean-field limit*. For brevity, we put the majority of derivations and proofs in a separate supplementary information note.

### Pathway-based generative model

Restricted-RNN is built upon a factor-centric view of neural computation which regards neurons as the substrate mediating the communication between factors (DePasquale et al., 2023). At the core of Restricted-RNN is the concept of **pathway** which mediates the communication between factors. Given a factor pair, the associated pathway consisted of one *input connectivity vector* and one *output connectivity vector*, of which the *input connectivity vector* embeds the sender factor information into neural population while the *output connectivity vector* reads out the embedded sender factor information from the neural population and sends it to the receiver factor. We adopted a subpopulation structure to construct both the *input connectivity vector* and the *output connectivity vector*.

Take the *k*_*i*_ → *k*_*j*_ pathway as an example. The associated *input connectivity vector* is a *MN* -dimensional vector (*M* is the number of subpopulations and *N* is the number of neurons in each subpopulation) given by:

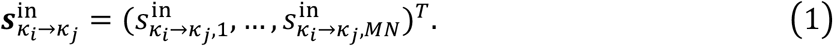

Here, each element 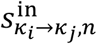 is generated through

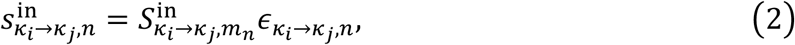

where *m*_*n*_ = ⌊(*n* − 1)/*N*⌋ + 1 denotes the index of the subpopulation that the *n*-th neuron belongs to (the first to the *N*-th neurons belong to subpopulation 1, the (*N* + 1)-th to the 2*N*-th neurons belong to subpopulation 2 and so on), 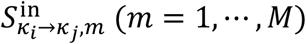 is the collective-level input connectivity strength, and 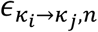 is the connectivity noise sampled from a standard normal distribution.

Similarly, the *output connectivity vector* for *k*_*i*_ → *k*_*j*_ is defined as:

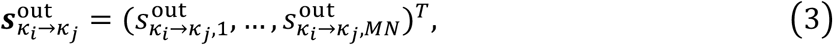

with each element 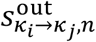 given by

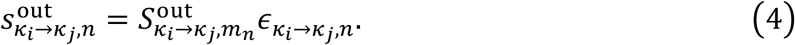

Here, 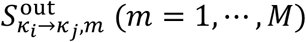 is the collective-level output connectivity strength and 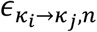 is the same Gaussian noise shared between the input and output connectivity vectors.

### The pathway-based factor graph and associated RNN

Now we can introduce factor graph—a graph modeling the complex interactions between factors through pathways. Consider a factor graph with *R* internal factors 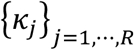 and *L* input factors 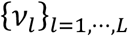. In total, it requires *R*^2^ + *LR* pathways, including 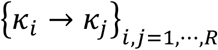 and 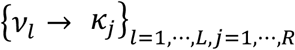, to model the arbitrary interactions between different factor pairs.

As each pathway can be expanded into the input and output connectivity of *MN* neurons, the factor graph corresponds to an RNN described by the following low-rank neural network (see Supplementary Note 1 for the derivation):

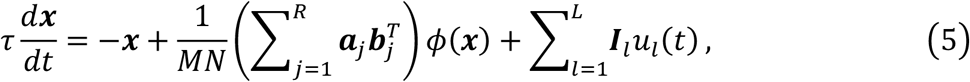

where ***x***(*t*) = (*x*_1_(*t*), ⋯, *x*_*MN*_(*t*))^*T*^ is the *MN*-dimensional vector (termed as hidden state) with each element representing the total input of the corresponding neuron, *ϕ*(·) is the nonlinear activation function, *u*_*l*_(*t*) is the *l*-th input signal, ***a***_*j*_ is a *MN*-dimensional connectivity vector with element

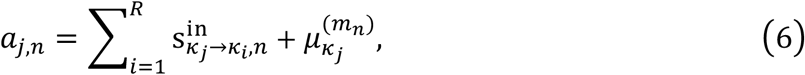

***b***_*j*_ is a *MN*-dimensional connectivity vector with element

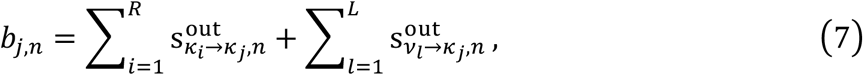

and ***I***_*l*_ is a *MN*-dimensional connectivity vector with element

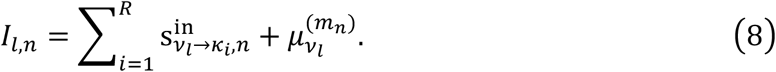

where 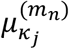 and 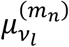 are parameters represents the subpopulation specific mean value for the corresponding task variable.

### The factor dynamics in the mean-field limit

While the RNN dynamics seems complex, the hidden state can be decomposed in the following way:

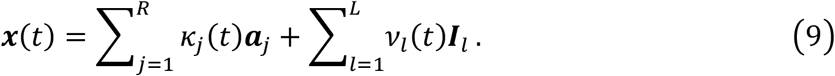

Importantly, in the mean-field limit (*N* → ∞), the dynamics of the internal factors *k*_*j*_(*t*) can be described by the following equation (see Supplementary Note 2 for the derivation):

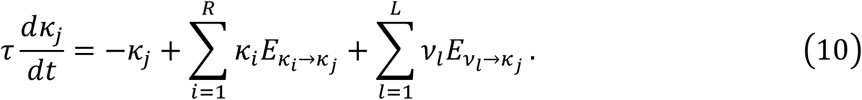

Here, *E*_*p*_ is the effective coupling strength of pathway *p* that can be written as

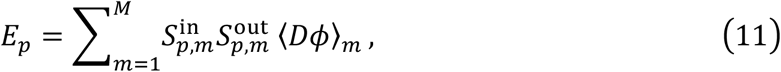

where ⟨*Dϕ*⟩_*m*_ is the average gain of the *m*-th subpopulation—a quantity depending on all factors, including *k*_*j*_(*t*) and *ν*_*l*_(*t*) (see Supplementary Note 2 for the details). Therefore, *E*_*p*_ is not a quantity with static value per se. Instead, it is dependent of not only the structural strength 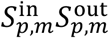 but also the state value *k*_*j*_(*t*) and *ν*_*l*_(*t*). Interestingly, when introducing the structural coupling vector

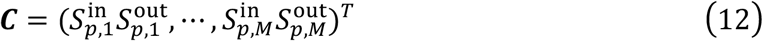

and

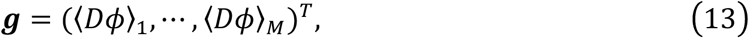

*E*_*p*_ can be written as the inner product between the structural coupling vector ***C*** and the subpopulation gain vector ***g***, a formula used in Fig. 5 to provide the geometric understanding for ubiquitous control in cognition.

### Details for modeling perceptual decision making (PDM) task (Figure 3)

#### Experiment details and electrophysiological recording (Fig. 3A, left)

A detailed description of the experiments can be found in Okazawa et al., 2021. The experiments involved a cohort of five adult macaque monkeys (Macaca mulatta) engaged in one of two perceptual decision-making tasks: motion direction discrimination and face discrimination.

##### Motion Direction Discrimination Task (motion task; Fig. 3A, top left)

Monkeys were trained to discriminate the overall direction of moving random-dots stimuli. The task difficulty was adjusted by altering the proportion of dots moving in a correct direction (coherence levels).

##### Face discrimination Task (face task Fig. 3A, bottom left)

Monkeys were trained to classify faces according to their species (monkey vs. human) or expression (happy vs. sad). A stimulus on each trial was sampled from a morph continuum between two prototype faces. The task difficulty was determined by the morph level; +100% or –100% morph corresponded to the two prototypes and 0% morph corresponded to the ambiguous, intermediate face stimulus.

The dataset included neural activity of 261 units recorded from the lateral intraparietal (LIP) area of the trained monkeys. 129 of them were recorded from three monkeys performing the motion task, while the remaining 132 units were recorded from the other two monkeys performing the face task. Spike counts were calculated from 100 ms before stimulus onset to 700 ms after stimulus onset, using non-overlapping 10ms bins.

#### Preference alignment and population average firing rates (Fig. 3A, right)

The neuronal activity of each unit was aligned according to its preferred saccade target (i.e., left or right target) following the procedure of Okazawa et al., 2021. The preference was determined using a separate memory-guided saccade task. Let *coh* denotes the signed stimulus evidence (e.g., in the motion task, *coh* represents motion coherence, its sign indicates the motion direction, and the absolute value indicates stimulus difficulty). For neuron *q, r*_*q*_(*coh, t*) is the average firing rate across all trials with the same *coh* at time *t* after stimulus onset. If the neuron’s preferred target corresponded to positive *coh*, the aligned response was defined as:

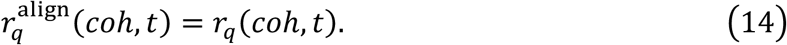

Otherwise, we flipped the neuron’s responses as:

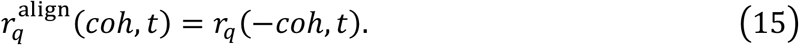

All firing rates reported in this section refer to the aligned neural activity and are denoted as

*r*_*q*_(*coh, t*) for simplicity. The population average firing rate was then computed by averaging across all recorded neurons:

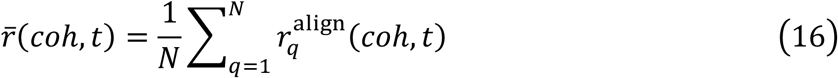

#### Determining the decision variable and stimulus difficulty axes in the neural state space using orthogonal canonical correlation analysis (Fig. 3, F-G; Fig. S3-5)

The reversal of mean firing rate 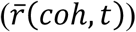 (Fig. 3A, bottom panel) indicates that neurons encode the stimulus difficulty (diff) in addition to the decision variable (dv). Here, we defined the dv as the signed stimulus evidence (*coh*) and the diff as the unsigned stimulus strength (|*coh*| ). Following the previous study (Okazawa et al., 2021), we identified the dv and diff axes in the neural state space using orthogonal canonical correlation analysis (CCA). For each trial *k*, we constructed a task parameter matrix *P* that

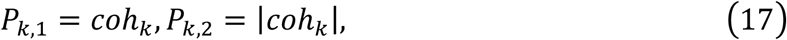

and a neural response matrix *R* of size *K* × *N*, where *R*_*k,n*_ was the mean firing rate of neuron *n* in a 100 ms window centered at 400 ms after stimulus onset. Neural response matrix was further mean-centered across trials before analysis. Orthogonal CCA was then applied to (*R, P*) to identify linear projections of neural activity that maximally correlated with dv and diff axes. Orthogonal CCA sequentially identified two pair of orthogonal projection vectors ***a***_*i*_ ∈ ℝ^*N*^, ***b***_*i*_ ∈ ℝ^2^, *i* = 1,2, such that the canonical variable *R****a***_*i*_ and *P****b***_*i*_ are maximally correlated. The resulting transformation matrices are *A* = [***a***_1_, ***a***_2_], *B* = [***b***_1_, ***b***_2_] and the neural encoding axes of dv (***β***^dv^) and diff (***β***^diff^) are given by the two columns of *AB*^−1^, which reflect the axis in the neural state space selective for each task variable.

### Determining the decision variable (dv) and stimulus difficulty (diff) axes in the neural state space using linear regression (Fig. S3-6)

To confirm that our results did not depend on CCA, we also estimated the dv and diff axes using linear regression. For the *i*-th neuron on trial *k*, let *r*_*i*_(*k*) denote the firing rates averaged over 350-450 ms time window after stimulus onset. It was expressed as a linear combination of task variables:

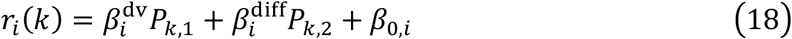

where *P*_*k*,1_ and *P*_*k*,2_ were the strength of dv and diff variables, respectively, and *β*_0,*i*_ was a bias term. To fit this linear model, we used Lasso regression with the regularization coefficient selected via five-fold cross-validation. The dv and diff axes were then given by 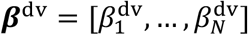 and 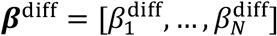.

#### Definition of three types of neurons (Fig. 3, C-E; Fig. S3-4)

Trials were partitioned by the stimulus conditions defined jointly by the evidence strength (i.e., easy or hard defined by the unsigned evidence strength) and the direction (i.e., the sign of the evidence *coh*) relative to the neuron’s preferred target. This resulted in four categories: non-preferred easy (NE, *coh* < −0.2), non-preferred hard (NH, −0.2 ≤ *coh* < 0), preferred hard (PH, 0 < *coh* ≤ 0.2) and preferred easy (PE, *coh* > 0.2). With Gaussian-smoothed (*σ*=15 ms) spike data, firing rates were computed in the 350−450 ms post-stimulus period using non-overlapping 10 ms bins. For each bin, the relative ordering of responses across the four conditions was evaluated, and a unit was assigned to a category only if the corresponding criterion was satisfied in all ten bins.

Neurons were considered dv-dominant if their firing rates increased monotonically with *coh* from the non-preferred easy to the preferred easy (Fig. 3C, top panel). Neurons were considered diff-dominant if responses in both hard conditions (non-preferred hard and preferred hard) exceeded responses in both easy conditions (Fig. 3C, bottom panel). Neurons that displayed a non-monotonic profile, e.g., with peak responses at intermediate evidence strengths but without meeting the strict difficulty-dominant criterion, were considered dv−diff mixed (Fig. 3C, middle panel). Neurons that did not meet any criterion were left unclassified. Proportions of each category in the model and monkey data are reported in Fig. S3-4, with corresponding example neurons shown in Fig. 3, C–E and Fig. S3-4, C–D.

#### ePAIRS analysis (Fig. 3, F-G; Fig. S3-6, A-B)

We used the ePAIRS (empirical Pairwise Angular Inference for Random Selectivity) analysis to assess whether neurons exhibited non-random clustering in the dv-diff selectivity coefficient space. The core idea of the method is that if neuronal selectivity coefficients are randomly distributed (e.g., independent samples from a multivariate Gaussian distribution), then the distribution of pairwise (between neurons) angles in high-dimensional space follows a well-defined null distribution. Significant deviations from this distribution indicate the presence of subpopulation structure.

Let *X* ∈ ℝ^*N*×*P*^ denote the matrix of selectivity coefficients, where each row corresponds to a neuron and each column to a task variable. Columns were mean-centered across neurons such that 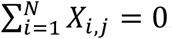. The *i*-th row of the matrix, denoted by ***x***_*i*_, represents the selectivity vector of neuron *i*. For each neuron pair (*i, j*), we compute the angle between selectivity vectors as

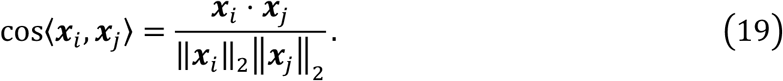

For each neuron *i*, the set of nearest neighbors *NN*_*k*_(*i*) was defined as the *k* neurons with the smallest angular distance from ***x***_*i*_. The empirical distribution of nearest-neighbor angles was then obtained by averaging over top-*k* neighbors for each neuron.

We generate the null distribution, using a multivariate Gaussian distribution that matches the covariance structure of the empirical data matrix *X*. From this distribution, surrogate datasets with the same number of neurons were sampled, on which the distribution of pairwise angles was computed as described above. This process was iterated 500 times to obtain a Monto Carlo estimate of the null distribution.

Finally, the empirical and null distributions were compared using a two-sided Wilcoxon rank-sum test to assess whether the empirical data can be explained by a single neuron population (i.e., whether the empirical distribution matches the null distribution).

#### Definition of simulated dataset for PDM task

There is a fixation period of duration *T*_fix_ = 100 *ms*, followed by a stimulus period of duration *T*_sti_ = 600 *ms* and then a delay period of duration *T*_delay_ = 500 *ms*. During stimulus period on trial *k*, it presents a stimulus-strength-dependent input 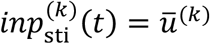, where 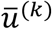 signifies stimulus strength supporting either choice 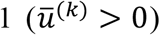 or choice 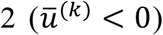. The stimulus-strength-independent input *inp*_const_(*t*) is always 1 during stimulus period and 0 during fixation and delay periods, regardless of stimulus strength.

During training, each batch includes 11 samples (trials) whose stimulus strengths are {−1, −0.5, −0.25, −0.125, −0.0625,0.0625,0.125,0.25,0.5,1.0,0} (denoted as trial 1 to trial 11).

The choice target *y*^(*k*)^ is defined as following: if 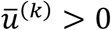, then *y*^(*k*)^ = 0; otherwise, *y*^(*k*)^ = 1. The choice target for trials with 0 stimulus strength is undefined, thus no constraints are placed on the model’s output for these trials.

The mean firing rates reversal puts additional constraints on the neuron activities. Therefore, we introduced representation targets 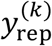 during training, which specify the desired aligned average firing rates (300−400 ms after stimulus onset) of all neurons during trials with stimulus strength of -1, 1, 0 (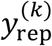, *k* = 1,10,11, respectively). The representation targets were set to (0, 0.2, 0.6) (corresponding to stimulus strength of -1, 1, 0, respectively) for motion task and set to (0, 0.9, 0.4) for face task.

#### Constructing and training Restricted-RNN for PDM task

In building a Restricted-RNN model for the PDM task, we run a systematic proposing-and-testing process for candidate hypotheses (specified by factor graphs). We progressively increased the model complexity until the minimal circuit capable of replicating the firing rate reversal was identified (Fig. 3B). We will detail the hypothesis models below.

##### Hypothesis 1 (H1): Single-factor circuit

H1 model consisted of one input factor *ν*_sti_ and one single factor (*k*_dv_). The network has only two pathways: *p*_1_ for input pathway from *ν*_sti_ to *k*_dv_, *p*_2_ for recurrent pathway of *k*_dv_. Each pathway is parameterized with two parameters 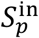 and 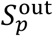, with *p* = *p*_1_, *p*_2_ . Because we only explore model with one subpopulation, the subpopulation index is omitted here. The input and output connectivity for each pathway is constructed as:

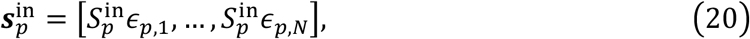

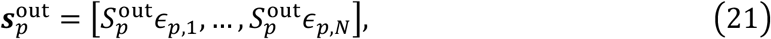

where the random variables *ϵ*_*p,i*_ are independently sampled from a standard normal distribution and shared across the input and output connectivity vectors of the same pathway. Mean components are included for all task variables.

The corresponding low-rank RNN is:

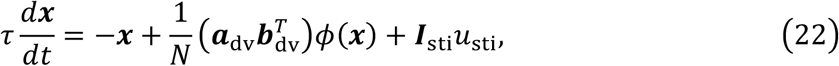

where

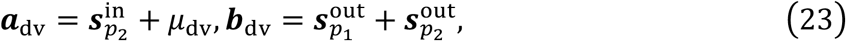

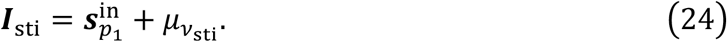

The network’s output is defined as:

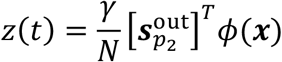

where *γ* is trainable parameter.

##### Hypothesis 2 (H2): Two-factor circuit

H2 model consisted of two input factors (*ν*_sti_ and *ν*_const_) and two single factors (*k*_dv_ and *k*_extra_). The network has four pathways: *p*_1_ for the input pathway from *ν*_sti_ to *k*_dv_, *p*_2_ for the recurrent pathway of *k*_dv_, *p*_3_ for the input pathway from *ν*_const_ to *k*_extra_ and *p*_4_ for the recurrent pathway of *k*_extra_ . Each pathway is parameterized with two parameters 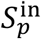 and 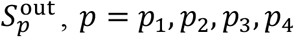.

Because we only explore model with one subpopulation, the subpopulation index is omitted here. The input and output connectivity for each pathway is constructed as:

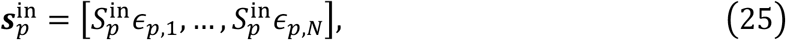

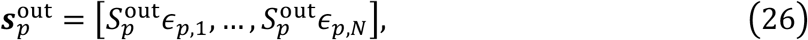

where the random variables *ϵ*_*p,i*_ are independently sampled from a standard normal distribution and shared across the input and output connectivity vectors of the same pathway. Mean components are included for all task variables.

The corresponding low-rank RNN is

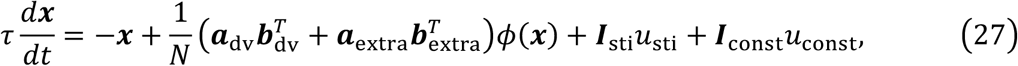

where

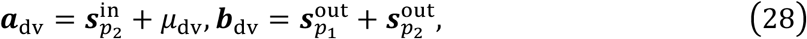

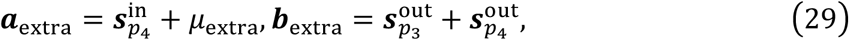

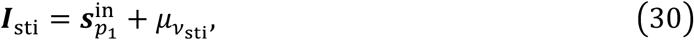

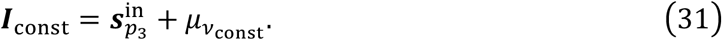

The network’s output is defined as:

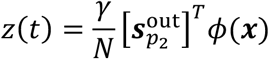

where *γ* is trainable parameter.

#### Training procedure and loss function

For both models (H1 or H2), 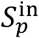 and 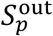 were initialized with normal distribution with zero mean and variance 0.01. Mean parameters (*μ*_·_) are initialized with 0. The RNN was discretized at 10 ms time steps with a time constant *τ* = 50 *ms*, and the activation function was the logistic sigmoid *ϕ*(*x*) = 1/(1 + *e*^−*x*^).

The Loss function includes two components:

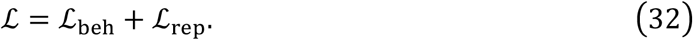

*L*_*beh*_ denotes the behavioral loss, aiming to optimize for correct choice, and is only applicable to trials with non-zero stimulus strength. It computes the cross-entropy between the model’s outputs and the trial targets in the delay period:

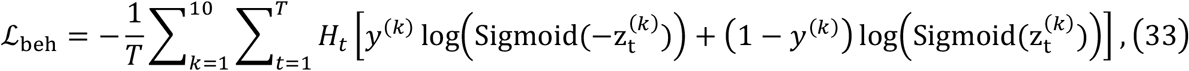

where *H*_*t*_ ∈ {0, 1} are masks (non-zero only during the delay period at the end of each trial).

The representational loss ℒ_rep_, a component providing constraints to the model representation, is calculated as the mean square error between representation targets 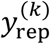 and the aligned average firing rates averaging over the period between time step 25 and 34 (10 steps in total, corresponding to 250-350 ms after stimulus onset):

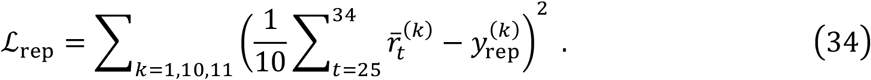

During testing, Gaussian white noise (*σ*_*inp*_ = 0.01) was added to all input factors. Model accuracy was defined as the proportion of trials (trial with zero-strength excluded) in which the sign of the network output at the final time step matched the stimulus strength. Models were considered successful if they achieved high behavioral performance (*L*_beh_ ≤ 0.02, and acc ≥ 0.99, over the final 100 batches) and satisfied representational constraints (*L*_rep_ ≤ 0.02 over the final 100 batches).

For the face task, only the two factors model (H2) satisfied the above criteria and was therefore selected as the final model. In this model, the second latent factor *k*_*extra*_ encoded stimulus difficulty and is henceforth termed *k*_diff_ in the following. We also observed that the mean parameters 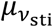 and 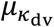 converged to values close to zero after training. Ablation tests confirmed that setting these parameters exactly to zero did not affect the model performance. We therefore fixed them zero in all following analysis for simplicity.

#### Explanation of why the H1 model produces a curved manifold but cannot reproduce the reversal phenomenon

For simplicity, we analyze the effect of the decision variable *k*_dv_ in the H1 model. We assume that *k*_dv_ increases monotonically with the stimulus strength *coh*, and that the neuronal activation function *ϕ*(·) is also monotonically increasing:

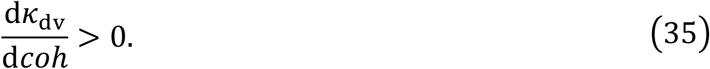

The neural activation vector is a linear embedding of the decision variable:

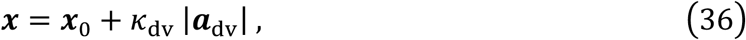

where |***a***_dv_| denotes the aligned dv axis. The observed neural activity corresponds to the element-wise nonlinear transformation of *x* through the activation function:

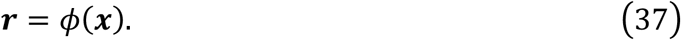

Although ***x*** lies on a straight line in the activation space, the nonlinear mapping *ϕ*(·) warps this line into a curved trajectory in the neural activity space. For example, if we let *k*_dv_ = *coh*, ***x*** = (*coh, coh* + 1), and *ϕ*(*x*) = max (*x*, 0) . Evaluating at *coh* = −1, 0, 1 yields ***r***|_*coh*=−1_ = (0,0), ***r***|_*coh*=0_ = (0,1), and ***r***|_*coh*=1_ = (1,2), which clearly do not lie on a straight line, illustrating how the curved manifold arises from the nonlinear transformation.

However, for each single neuron *i*, the activity *ϕ*(*x*_*i*_) remains a monotonic function of the stimulus strength:

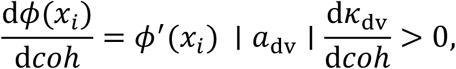

because both *ϕ*^′^(*x* ) and 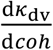 are positive by assumption. Therefore, each neuron’s firing rate increases monotonically with stimulus strength, and no mean-rate reversal can occur. In other words, while the element-wise nonlinearity can bend the population trajectory to create a curved manifold, it cannot invert the monotonic relationship between stimulus strength and firing rate. Reproducing reversal phenomena thus requires additional mechanisms beyond a single monotonic nonlinear transformation, such as stimulus difficulty representation.

#### Collective dynamics of Restricted-RNN in PDM task (Fig. S5-2)

The two internal factors *k*_dv_ and *k*_diff_ have the following dynamics:

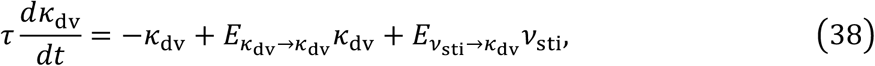

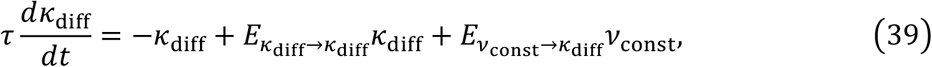

where *E*_*p*_ denotes the effective coupling of pathway *p*, given by:

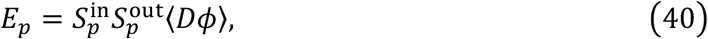

where ⟨*Dϕ*⟩ is the subpopulation gain given by:

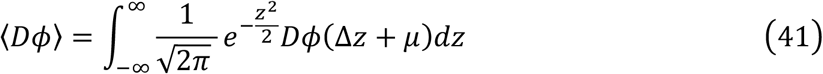

with

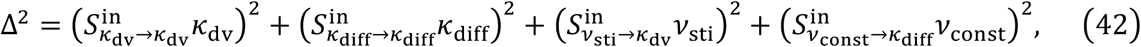

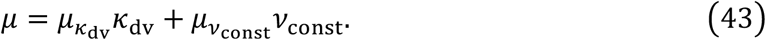

Although *ν*_const_ is independent of stimulus strength, the subpopulation gain depends on it. Therefore, the information flow from *ν*_const_ to *k*_diff_ varies with the strength of the stimulus. Our numerical simulations indicate that as strength increases, the subpopulation gain first increases and then decreases, with the maximum value corresponding to zero stimulus strength (Fig. S3-7A).

### Details for modeling sequence working memory (SWM) task (Figure 4)

#### Experiment details and electrophysiological recording

A detailed description of the experiment can be found in Chen et al., 2024. Briefly, two macaque monkeys were trained to perform a sequence working memory task. Each trial began with a fixation period, after which the monkeys were sequentially presented with 1-3 visual stimuli which are white dots at distinct locations chosen from six candidates (labeled 1 to 6, lasting for 250 ms), with a randomized inter-stimulus interval of 300-500 ms. This is referred to as the sensory period. The monkeys were required to maintain the stimuli in memory during the delay period and then report the presented locations in the order they appeared.

While neural activity was recorded simultaneously from multiple brain regions, only recordings from the frontal cortex were analyzed in the present study. We focused on length-3 correct trials. This dataset comprised 3,225 units (monkey 1: 2,058 units; monkey 2: 1,167 units). Monkey 1 and monkey 2 correspond to monkeys O and G, respectively, as in the previous study.

Neural activities were converted into spike counts using 100 ms bins with 50 ms steps (50 ms overlap between consecutive bins). Each trial was aligned to four task-defined periods: (1) a window aligned to the onset of the first stimulus and lasting 650 ms (12 bins); (2) a window aligned to the onset of the second stimulus and lasting 650 ms (12 bins); (3) a window aligned to the onset of the third stimulus and lasting 1600 ms (31 bins, covering both the third stimulus and early delay); and (4) a late delay window defined as the final 600 ms preceding the go cue (11 bins). In total, each trial consisted of 66 time bins.

#### Linear regression and tuning curve

To quantify how target locations affect the neural activity in SWM task, we fit a multivariate linear regression model to each neuron. The firing rate of the *i*-th neuron at time *t* on trial *k* (denoted as *y*_*i,t*_(*k*)) was expressed as following:

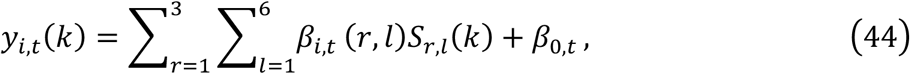

where *r* refers to the ordinal rank of element in the sequence and *l* denotes the index of location. *S*_*r,l*_(*k*) is an indicator variable for target locations that equals 1 if the rank *r* of trial *k* was location *l*, and 0 otherwise. For example, for the sequence [5,3,1], the active regressors would be *S*_1,5_, *S*_2,3_ and *S*_3,1_, while all the others would be zero. To prevent overfitting, we used Lasso regression with a regularization coefficient selected by 5-fold cross-validation. For each neuron *i* and rank *r* at time *t*, we obtained six regression coefficients {*β*_*i,t*_(*r*, 1), …, *β*_*i,t*_(*r*, 6)}, corresponding to the six possible locations. These coefficients defined the neuron’s tuning curve for locations of rank *r* at time *t*.

The tuning curve for the rank *r* during the sensory period was defined as 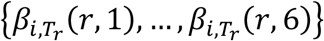 where subscript *T*_*r*_ refers to the 100-400 ms window after the *r*-th stimulus onset. In computing these coefficients, the target 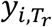 corresponds to the firing rates averaged over *T*_*r*_ (Fig. 4H). Similarly, the tuning curve for rank *r* at the late delay period was defined as 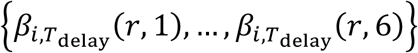 with *T*_de*l*ay_ being the 600 ms window before the go cue.

#### Estimating activation amplitude and tuning phase (Fig. 4H-K; Fig. S4-3; Fig. S4-4; Fig. S4-5; Fig. S4-6)

We quantified the tunning curve by fitting it with a cosine function. First, the six locations were mapped to angles 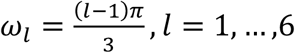, between 0 and 2*π*. Then we fit the tuning curve with

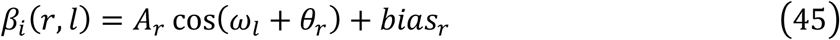

where *A*_*r*_ denotes the **activation amplitude**, *θ*_*r*_ is the **tuning phase**, and *bias*_*r*_ is a baseline offset. Fitting was performed using a standard nonlinear least-squares solver with constraints *A*_*r*_ ≥ 0 and *θ*_*r*_ ∈ [0,2*π*) (Figs. S4-4, A and B).

For each neuron, the activation amplitudes for the three ranks were summarized by triplets 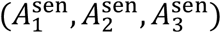 for the sensory period and 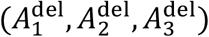 for the late delay period, and the tunning phases by 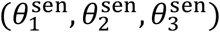 and 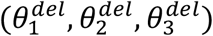. The amplitude triplets and the tuning phase triplets provide a comprehensive measure of a neuron’s response pattern to stimuli, forming the basis for subsequent analysis.

#### Normalized participation ratio and variability across rank (Figs. 4, J and K; Fig. S4-4, C-K; Fig. S4-5; Fig. S4-6)

The neuron *i*’s contribution to the representation of the rank-*r* sensory input was measured by 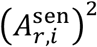. To identify the proportion of neurons that contributed substantially, we computed the normalized participation ratio (NPR):

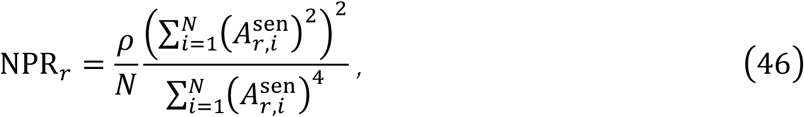

where *ρ* = 1 by default and *N* is the number of all neurons. Intuitively, NPR estimates the fraction of neurons making significant contributions to the *r*-th sensory input. For example, an NPR of 0.1 indicates that the neurons with the largest 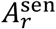 (top 10%) values were considered as significantly responding to the *r*-th sensory input. Only neurons that significantly responded to at least one rank were included in subsequent analysis, with the number being denoted as *N*_*ρ*_.

We next asked whether rank modulated each neuron’s representation of spatial location during the sensory period. Specifically, we examined whether the preferred location was consistent across the three ranks (equivalently, whether the tuning phase shifted with rank). To quantify this, we investigate the variability in both tuning phases and amplitudes across ranks.

##### Phase variability

We use two indices to capture the variability of tunning phases for different rank: (1) phase difference (*θ*_diff_) is the mean pairwise difference among tuning phases of different ranks:

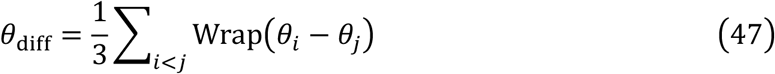

where Wrap(·) maps values to [0, *π*] (Fig. S4-4, C-H); (2) phase variability *θ*_std_ is the circular standard deviation of (*θ*_1_, *θ*_2_, *θ*_3_) (Fig. S4-4, I-K).

##### Amplitude variability

To visualize whether amplitudes differed across ranks, we plotted the triplet 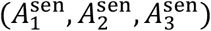 in a triangular coordinate system, in which the ratio of a point’s distances to the three sides is equal to the amplitude ratio 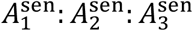. In this way, the three vertices indicate pure selectivity for one rank, while the center point corresponds to equal amplitudes across all ranks 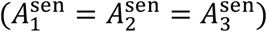. In the experimental data, many neurons were located away from both the center and the vertices, indicating mixed selectivity rather than either pure or uniform tuning (Fig. 4, J and K). To ensure the robustness of the conclusion, we also varied the coefficient *ρ* to include different number of neurons (Fig. S4-6). The results remained consistent under these conditions.

#### Estimate the number of clusters (Fig. 4, J and K; Fig. S4-5A; Fig. S4-6)

In Figure 4K, the neuron distribution in amplitude triangular space demonstrates clear clustering pattern, with many neurons concentrated near three distinct regions. To estimate the number of clusters, we fit Gaussian mixture models (GMM) to the distribution of the amplitude triplets.

We first gathered the tuning amplitude of neurons across ranks to form a matrix *A* ∈ ℝ^*N*×3^, where each row corresponding to a triplet 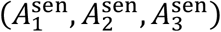 for the corresponding neuron. We considered GMMs with *K* = 1, …,5 components. For each *K*, we fit the model using half-split cross-validation for *R* = 1000 repetitions: each time, neurons were randomly partitioned into 50% training and 50% test sets. Then the GMM was fit to the training set using expectation-maximization algorithm, and the mean log-likelihood on the test set was computed. This procedure yields a cross-validated log-likelihood matrix (denoted as *LL*) of shape 1000 × 5.

The optimal cluster number 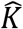 was defined as the *K* that maximizes the median cross-validated log-likelihood. To assess whether 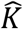 significantly outperformed alternatives, we applied one-sided Wilcoxon signed-rank test on the log-likelihood difference 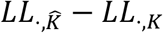. The resulting *p*-values were FDR corrected for multiple comparisons (Fig. 4K, right panel for monkey 1; Fig. S4-5, A, right for monkey 2; Fig. S4-6, A and B, middle for monkey 1 with scaled NPR; Fig. S4-6, C and D, middle for monkey 2 with scaled NPR). For each cross-validation repetition, we also recorded *K*_*max*_ achieved the highest held-out log-likelihood. The resulting frequency distribution of *K*_*max*_ across 1,000 repetitions provided a measure of the robustness of the log-likelihood based clustering analysis (Fig. S4-5B for monkey 1; Fig. S4-5C for monkey 2; Fig. S4-6, A and B, right for monkey 1 with scaled NPR; Fig. S4-6, C and D, right for monkey 2 with scaled NPR).

#### Definition of simulated dataset for SWM task

A length-*L* trial was defined as follows. A trial began with a 200 ms fixation period, followed by *L* stimulus each lasting for 200 ms, separated by 400-600 ms inter-stimulus intervals. Then the trial ends with a 1000 ms delay period. In the *k*-th trial, the stimuli 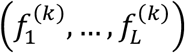 were randomly drawn (with replacement) from the set {1, …,6}.

Three input channels are assumed for the simulation. Stimulus locations were encoded in the first two input channels, *u*_*x*_ and *u*_*y*_, corresponding to the *x*- and *y*-coordinates on a unit circle. Each stimulus index *f*_*i*_ was mapped to a fixed location on the circle 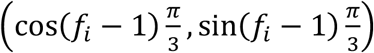.

During the rank-*r* stimulus presentation, the inputs were set to the corresponding coordinates,

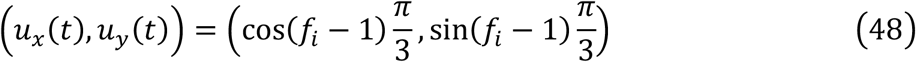

and set to zero during all other periods. The third channel encodes the location-independent input *u*_*const*_(*t*) indicating the presence of a stimulus: it equals 1 during any stimulus presentation, and 0 at all other times. The model was trained to reproduce the original stimulus sequence 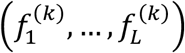. Therefore, the target output for each trial was a (2*L*)-dimension vector:

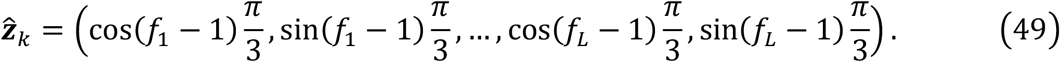

#### Constructing and training Restricted-RNN for SWM task

The Restricted-RNN for SWM task incorporated three input factors (*ν*_*x*_, *ν*_*y*_ and *ν*_*const*_) and seven internal factors (*k*_1,*x*_, *k*_1,*y*_, *k*_2,*x*_, *k*_2,*y*_, *k*_3,*x*_, *k*_3,*y*_, *k*_*c*_) . Note that in the main text, we define the stimulus input factor *ν*_*sti*_ = [*ν*_*x*_, *ν*_*y*_] and rank-*r* working memory factor *k*_*r*_ = [*k*_*r,x*_, *k*_*r,y*_]. For brevity, here we expand these factors into x- and y-components. The network has 14 pathways: *p*_*r,x*_ for input pathway from *ν*_*x*_ to *k*_*r,x*_ (*p*_*r,y*_ for input pathway from *ν*_*y*_ to *k*_*r,y*_), *r* = 1,2,3, *p*_*r*+3,*x*_ for recurrent pathways of *k*_*r,x*_ (*p*_*r*+3,*y*_ for *k*_*r,y*_), *p*_7_ for input pathwaty from *ν*_*const*_ to *k*_*c*_, and *p*_8_ for recurrent pathway of *k*_*c*_. Each pathway *p*_*r,x*/*y*_ was parameterized by subpopulation-specified scalars 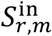 and 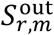 (same parameter for *x* and *y, r* = 1, …, 6), with *m* = 1, …, *M*, where *M* is the number of subpopulations. Pathway *p*_7_ and *p*_8_ was parameterized by subpopulation-specified scalars 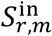 and 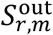, with *r* = 7,8 and *m* = 1, …, *M* (See Fig. S4-2).

For an instantiated low-rank RNN, assuming each subpopulation contains *N* neurons, the **input and output connectivity vector** for pathway *p* are constructed as:

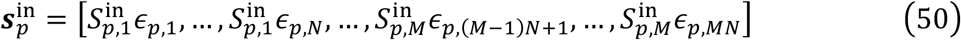

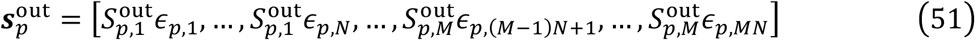

where the random variables *ϵ*_*p,i*_, *i* = 1, …, *MN* are independently sampled from a standard normal distribution and shared across the input and output connectivity vectors of the same pathway. Mean component was included for the biased input and the internal control factor *k*_*c*_. Combining the connectivity vectors across all pathways yields the corresponding low-rank RNN (see Fig. S4-2C), with the following dynamics:

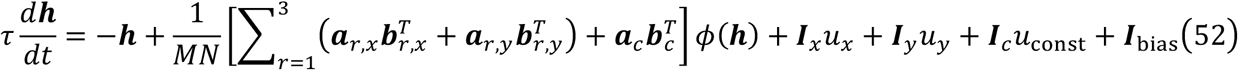

with

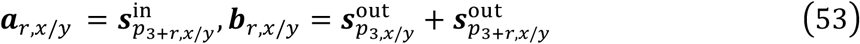

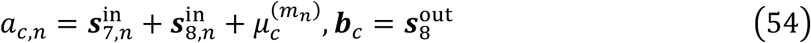

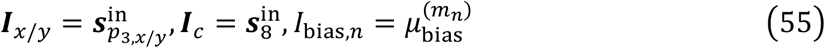

The model has six output channels ([*z*_*r,x*_, *z*_*r,y*_] for rank-*r* memory readout) defined as

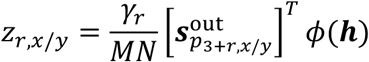

where *γ*_*r*_, *r* = 1,2,3 are trainable parameters controlling the readout strength. The activation function is *softplus* function defined as:

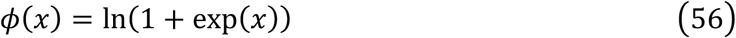

#### Loss function and training procedure

The loss function is composed of three parts:

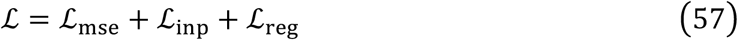

the term ℒ_*mse*_ is the mean-square error between network’s output and the target output:

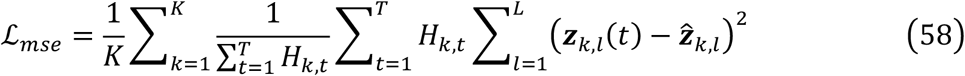

where *T* is the number of time steps, *H*_*k,t*_ is 0-1 mask that equals 1 during the delay period and 0 otherwise.

The ℒ_*inp*_ implements the input pathway gating hypothesis (Fig. 4C) by directly constraining the temporal profile of input pathway effective coupling strength:

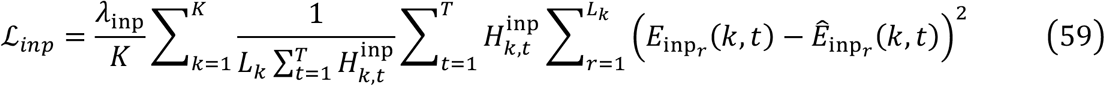

where *L*_*k*_ is the length of the *k*-th trials, 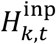 is 0-1 mask that equals 1 during any stimulus and 0 otherwise, and *λ*_inp_ = 1. In this equation, 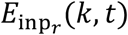 is the effective coupling of rank-*r* input pathway at time *t*:

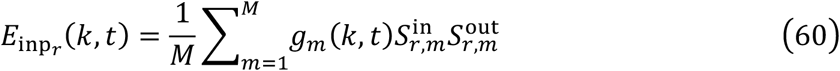

where *g*_*m*_(*k, t*) is the *m*-th subpopulation gain:

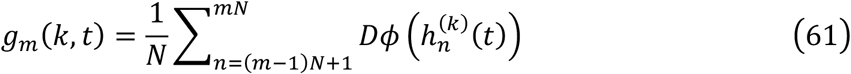

where 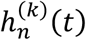 denoted the activation of the *n*-th neuron at time *t* during trial *k*. And 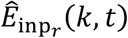 is the target effective coupling strength was set to 1 during the rank-*r* stimulus and 0 otherwise. The term ℒ_*reg*_ is the L2 regularization term defined as:

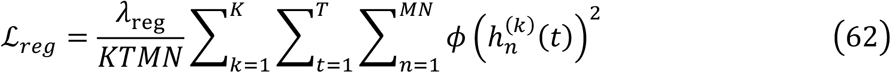

Training followed a staged curriculum. The first stage lasts for 1000 epochs, the second last for 5000 epochs and the third last for 45,000 epochs. The coefficient *λ*_re*g*_ is set to 10^−5^ during the first 46,000 epochs and 0.01 during the last 5,000 epochs. At the L stage, the model was trained on length 1 to L dataset. The length-*L* dataset enumerated all 6^*L*^ sequence. 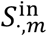 and 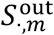 are initialized with normal distribution with zero mean and standard deviation of 0.01, mean components are initialized with zero and *γ*_*r*_ are initialized with 1. The training is performed using the PyTorch framework with the Adam optimizer with a learning rate of 0.001.

#### Collective dynamics of Restricted-RNN in SWM task (Fig. S5-1)

The internal factors in this task have the following dynamics:

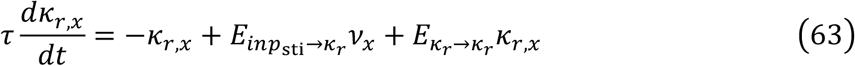

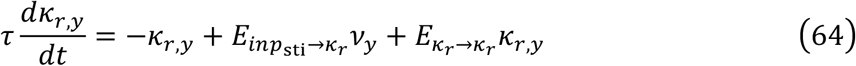

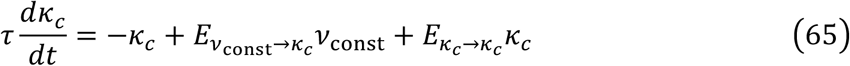

Because the two pathways *ν*_*x*_ → *k*_*r,x*_ and *ν*_*y*_ → *k*_*r,y*_ share the same effective coupling profile, we use a single notation 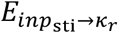 for both pathways. Likewise, 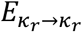 denote the effective coupling of the recurrent pathways for *k*_*r,x*_ and *k*_*r,y*_.

The effective coupling of each pathway is the gain weighted sum over subpopulations:

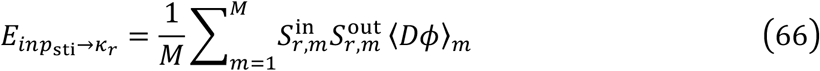

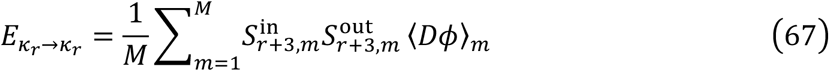

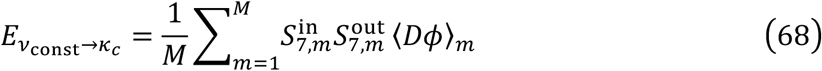

Where the subpopulation gain in the mean-field form is:

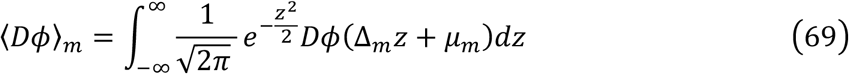

with

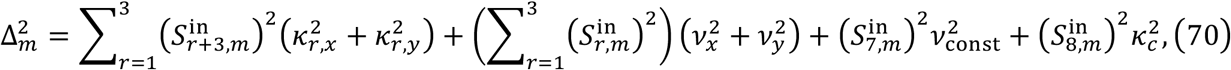

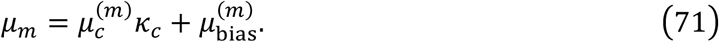

#### Mechanism of stimulus-gated input

Although 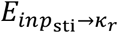 depends on all factors via ⟨*Dϕ*⟩_*m*_, the internal factor *k*_*c*_ is the primary modulator of the gating profile. As successive stimuli arrive (rank 1 to 3), *k*_*c*_ increases in steps (empirically near 0 before rank-1, ∼0.5 before rank-2, and 1 before rank-3), which shift *μ*_*m*_ so that 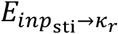 is active only during the corresponding stimulus window and suppressed otherwise, realizing the proposed input pathway gating mechanism.

## Supplementary figure

**Fig. S3-1.**
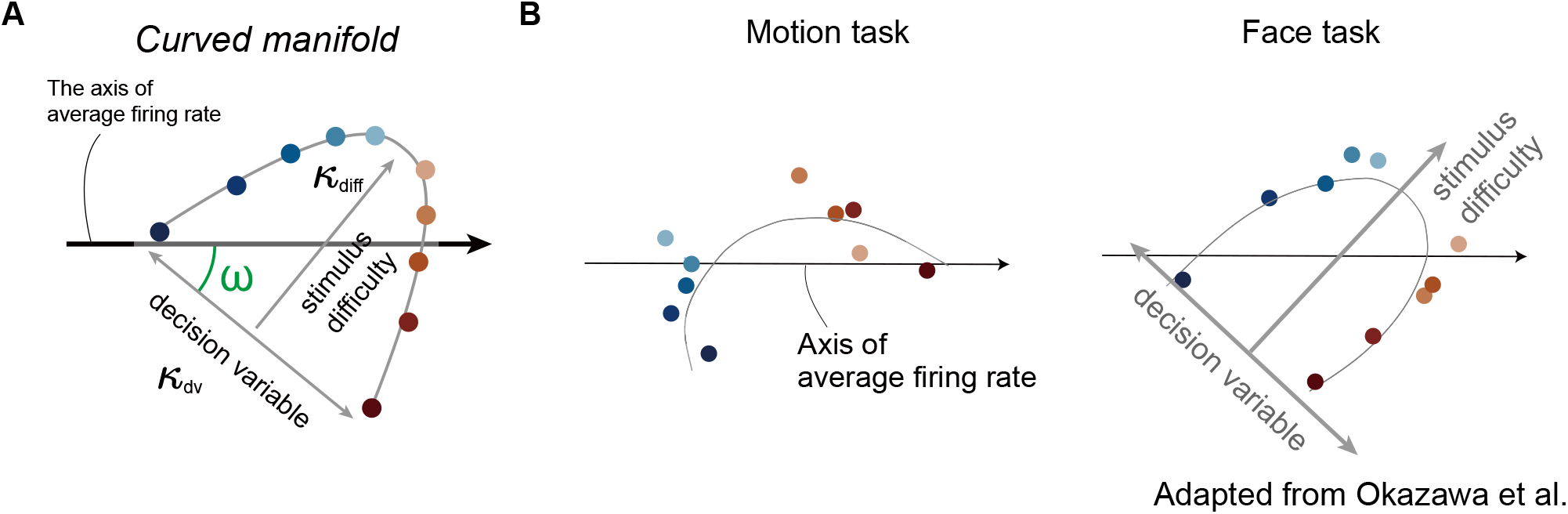
Curved manifolds in the motion and face tasks. (A) Schematics of curved manifold. (B) Curved manifolds in experimental data for the motion (left) and face (right) tasks (adapted from Okazawa et al., 2021).

**Fig. S3-2.**
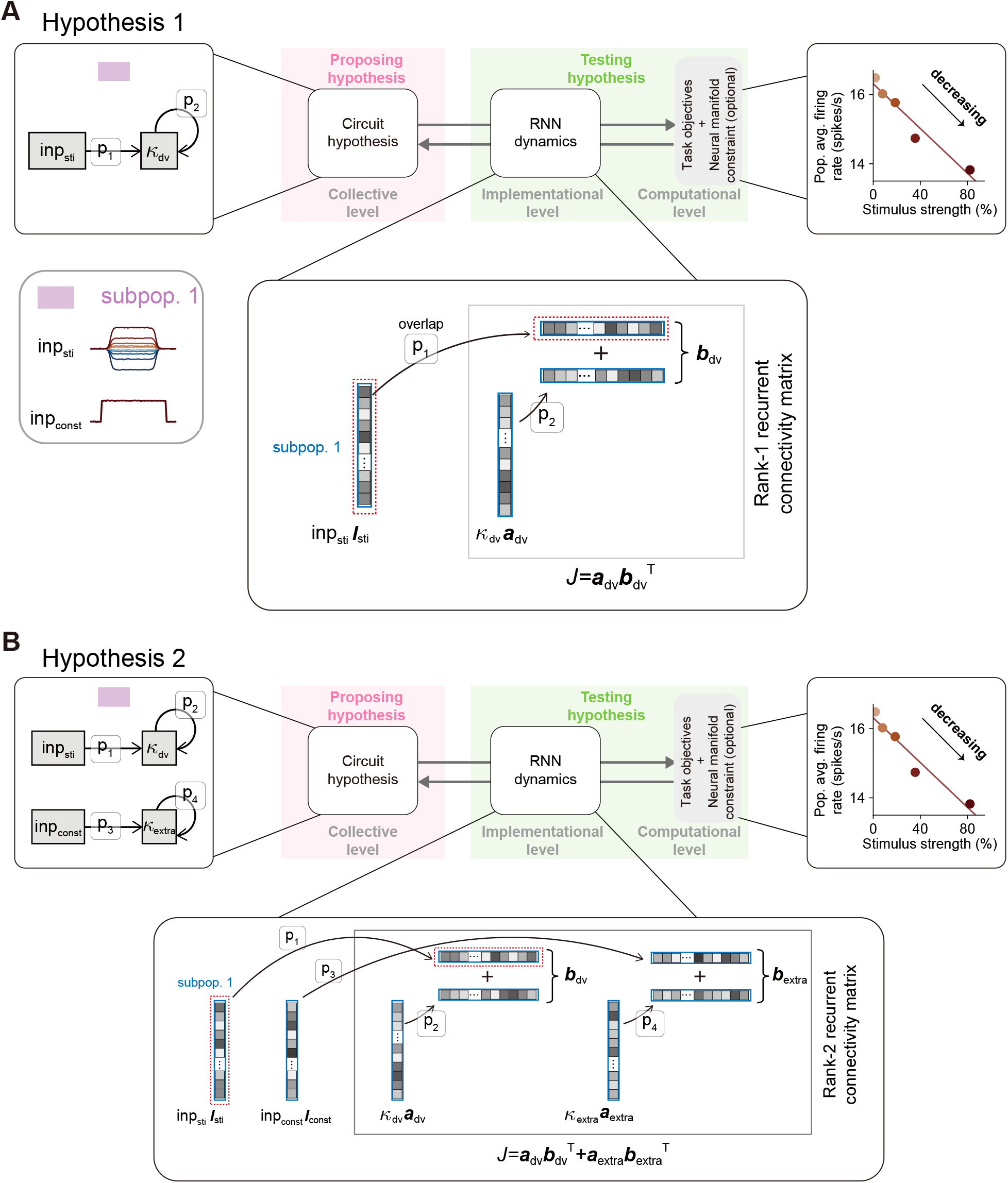
Hypothetic architectures of possible Restricted-RNN models for the PDM task. (A and B) A summary of applying the Restricted-RNN training framework to the PDM task for the hypothesis 1 (A) and hypothesis 2 (B). The task objective is to reproduce the mean firing rate reversal, and our results indicate that only hypothesis 2 successfully achieves this.

**Fig. S3-3.**
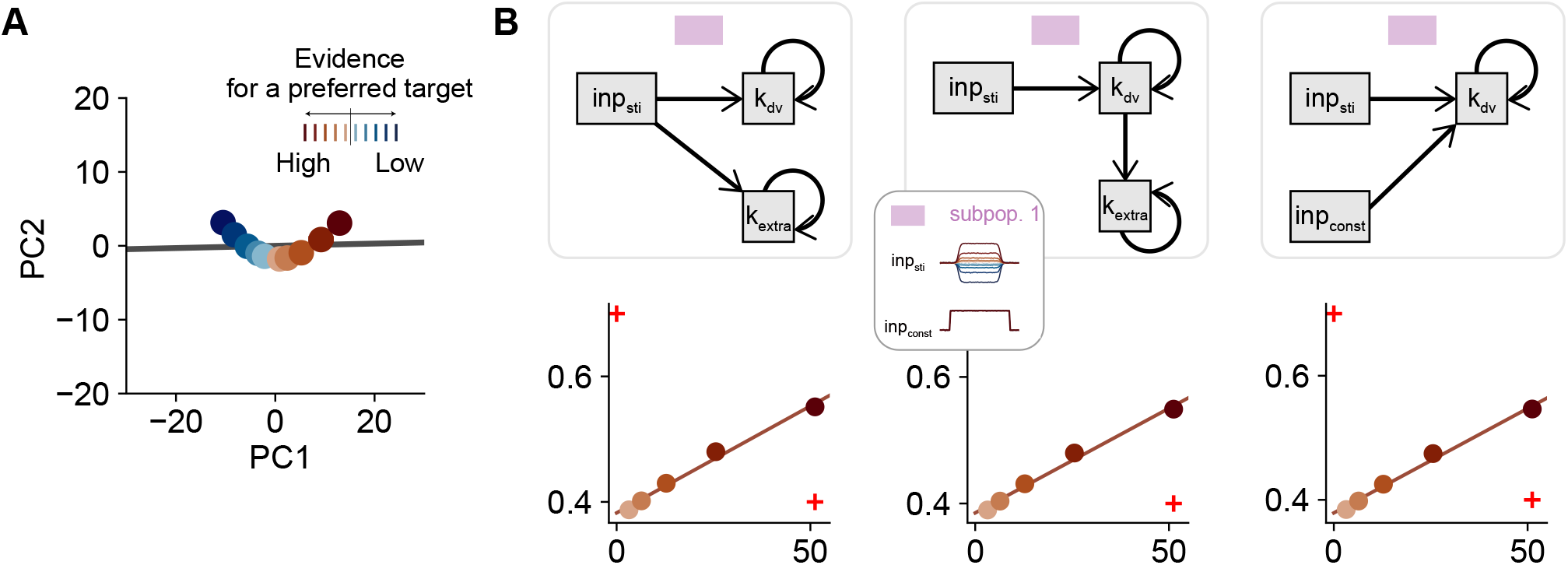
Behavior of the H1 Restricted-RNN and its variants for the PDM task. (A) Curved manifold of the model for the H1 hypothesis. (B) The collective circuits of three additional hypotheses (upper) and their corresponding mean firing rates curves (bottom). All these hypotheses fail to generate mean firing rates reversal. The nearly identical mean firing rate curves of the three models show that, unlike in H2, *k*_*extra*_ and *inp*_*const*_ do not affect the models’ behavior.

**Fig. S3-4.**
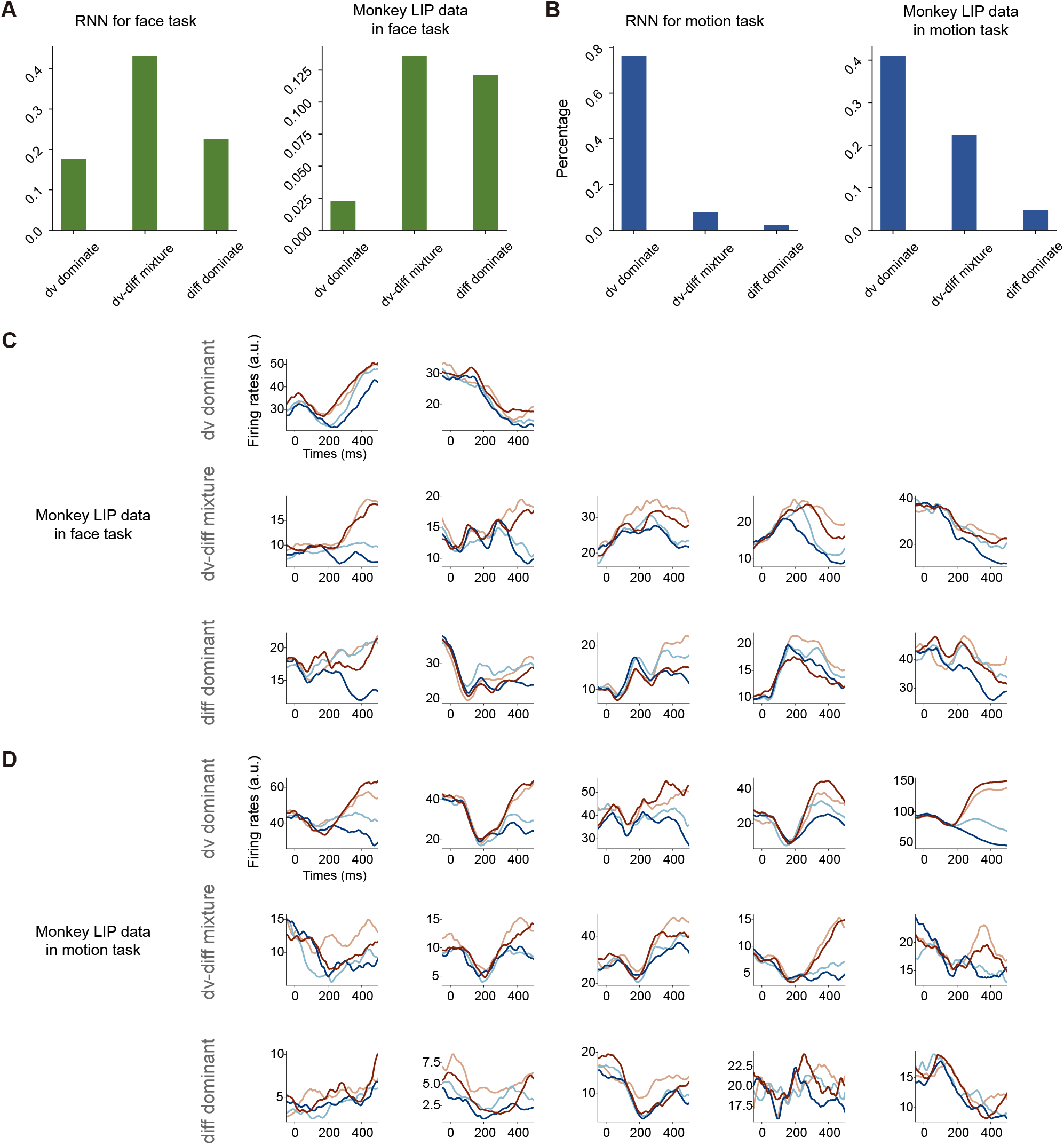
Proportion of neurons of different types and corresponding example neurons from the Restricted-RNN model and monkey data for the PDM task. (A) Proportion of different types of neurons in the model (left) and monkey data (right) for the face task. In the model, there are 17.8% dv-dominant neurons, 43.3% dv-diff-mixed neurons, and 22.5% diff-dominant neurons. In the monkey data, there are 3 (2.3%) dv-dominant neurons, 18 (13.6%) dv-diff-mixed neurons, and 16 (12.1%) diff-dominant neurons. (B) Proportion of different types of neurons in the model (left) and monkey data (right) for the motion task. In the model, there are 76.8% dv-dominant neurons, 7.8% dv-diff-mixed neurons, and 2.4% diff-dominant neurons. In the monkey data, there are 53 (41.1%) dv-dominant neurons, 29 (22.5%) dv-diff-mixed neurons, and 6 (4.7%) diff-dominant neurons. (C) Example neurons of different types in the monkey data for the face task. (D) Example neurons of different types in the monkey data for the motion task.

**Fig. S3-5.**
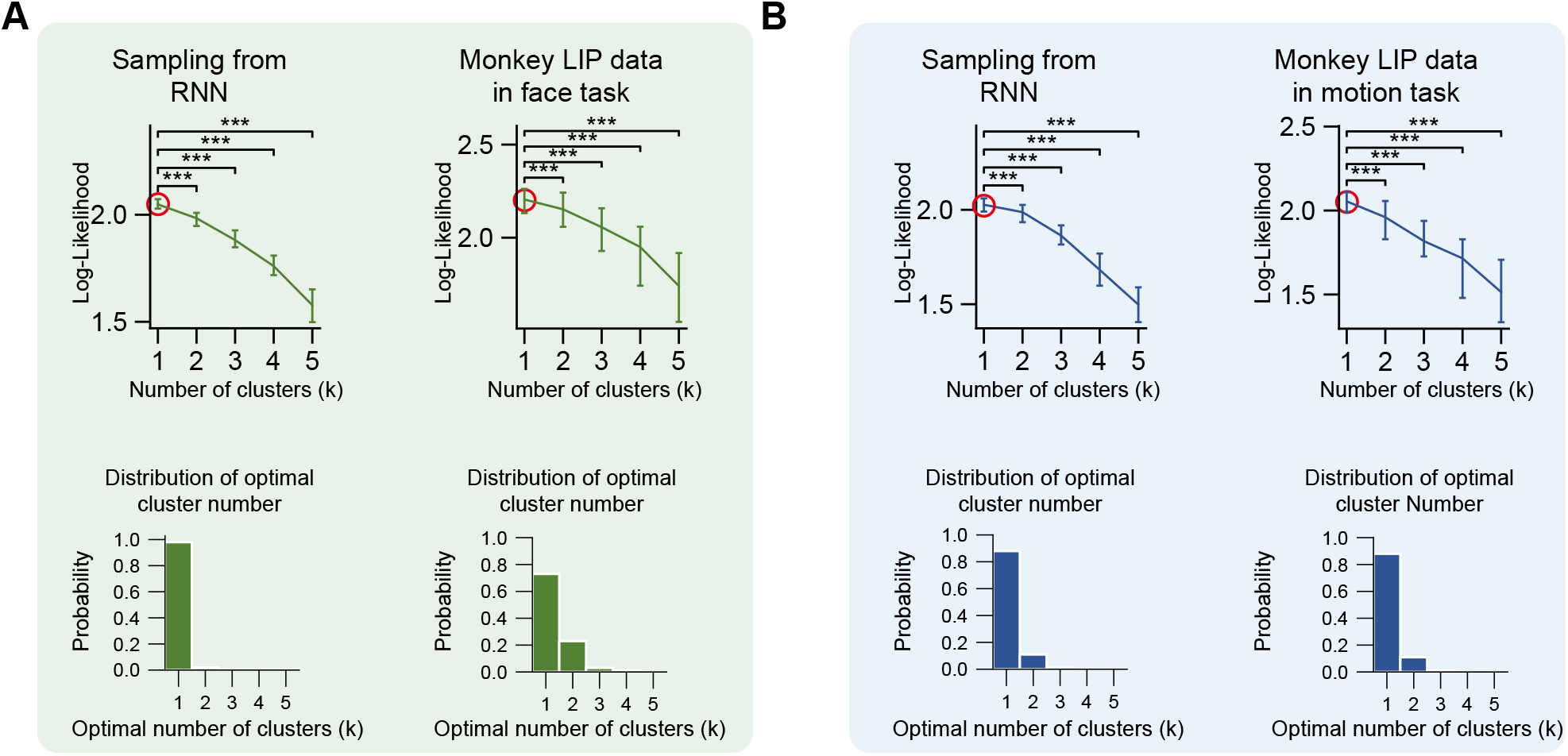
Clustering analysis in the space defined by the dv and diff axes from CCA. (A) Comparison of the cluster structures of our RNN model and monkey LIP data in face task. Likelihood-based clustering analysis (bootstrapped; error bars show s.e.m.) suggests one cluster for both cases (upper row). The distribution of optimal cluster numbers (bottom row) supports the same conclusion. (B) Same as (A) but for motion task. Again, one cluster is identified in both the RNN model and monkey LIP data.

**Fig. S3-6.**
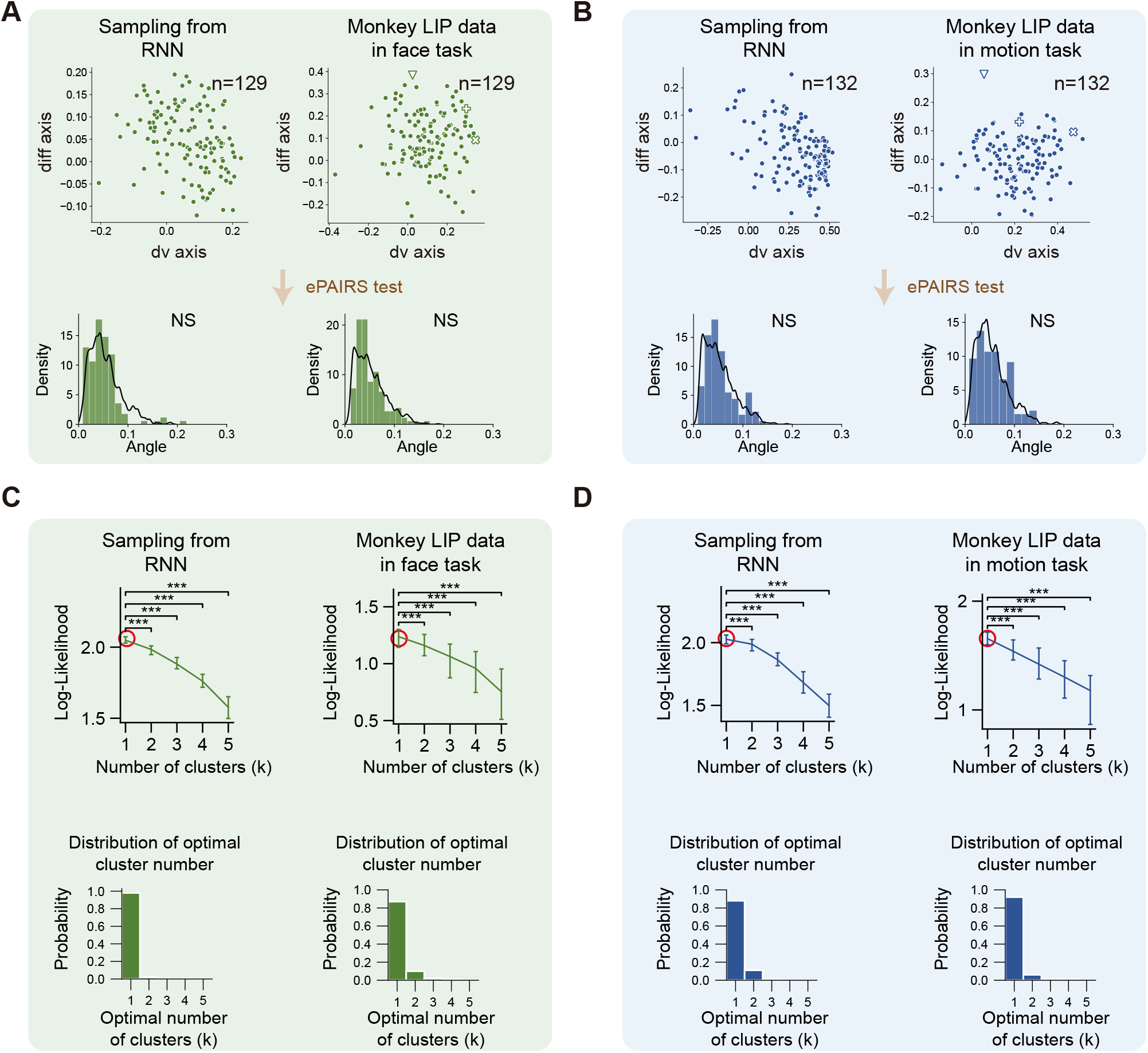
Clustering analysis in the space defined by the dv and diff axes from linear regression. (A) Analysis of population structure in the dv-diff space for face task. Each point represents the activity of a neuron projected to this space. ePAIRS test suggests single population for both the Restricted-RNN model and the monkey LIP data. (B) Same as (A) but for motion task. ePAIRS test also suggests single population for both the Restricted-RNN model and the monkey LIP data. (C) Comparison of the cluster structures of our RNN model and monkey LIP data in face task. Likelihood-based clustering analysis (bootstrapped; error bars show s.e.m.) suggests one cluster for both cases (upper row). The distribution of optimal cluster numbers (bottom row) supports the same conclusion. (D) Same as (C) but for motion task. Again, one cluster is identified in both the RNN model and monkey LIP data.

**Fig S4-1.**
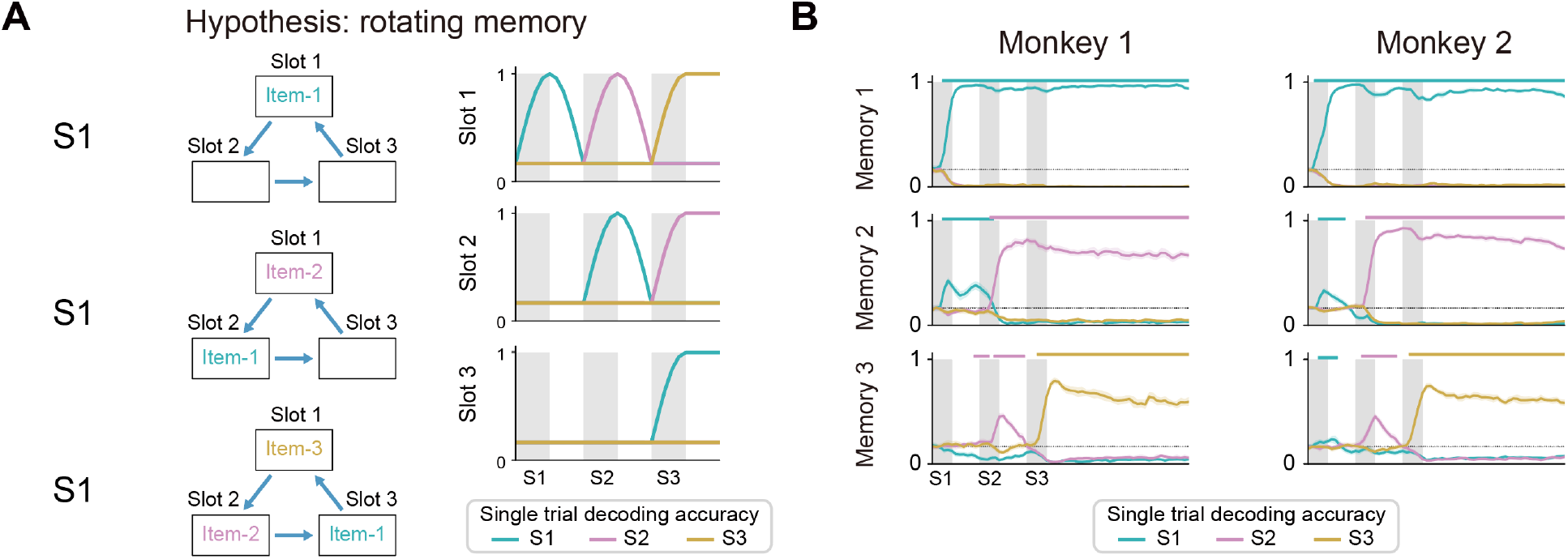
The rotating memory hypothesis for the SWM task is inconsistent with experimental data. (A) Schematics of the rotating memory hypothesis (adapted from Whittington et al., 2024). (left) The sensory input always enters slot1. Upon each input, previous items move to the subsequent slots (e.g., slot1 to slot2, slot2 to slot 3). (right) Predicted decoding accuracy in each slot at different sensory input periods. (B) Data from both monkeys are inconsistent with the rotating memory hypothesis. (Similar results have been found in Chen et al., 2024). The sensory inputs at different ordinal ranks directly enter into corresponding rank subspaces without rotating.

**Fig S4-2.**
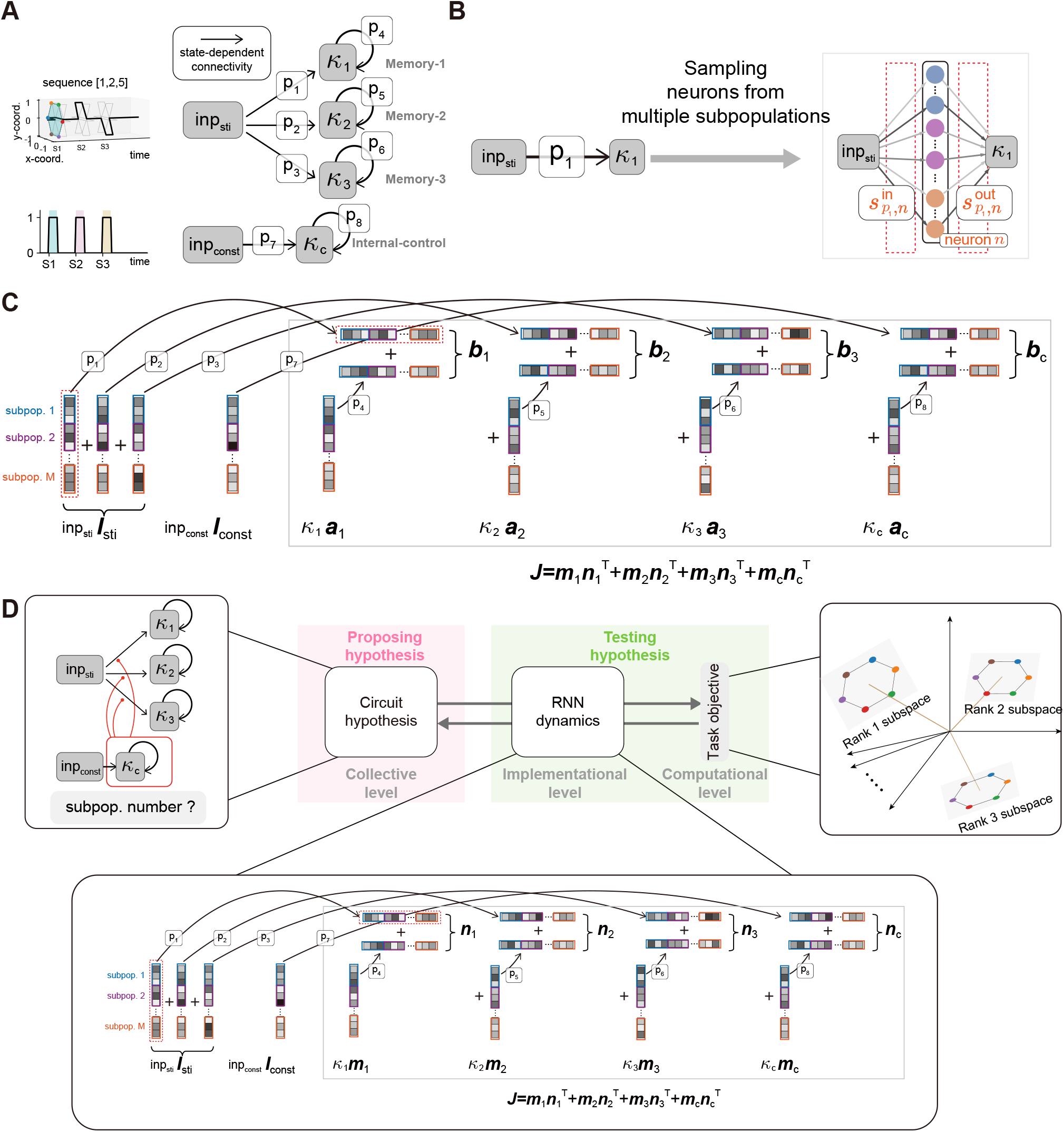
Architecture of the Restricted-RNN model for the SWM task. (A) Collective-level circuit of our Restricted RNN model for the SWM task. (B and C) Constructing low-rank RNN (C) from the collective circuit through the pathway-based generative model (B) (see Fig. S2-3 for a general description). (D) A summary of applying the Restricted-RNN training framework to the SWM task.

**Fig S4-3.**
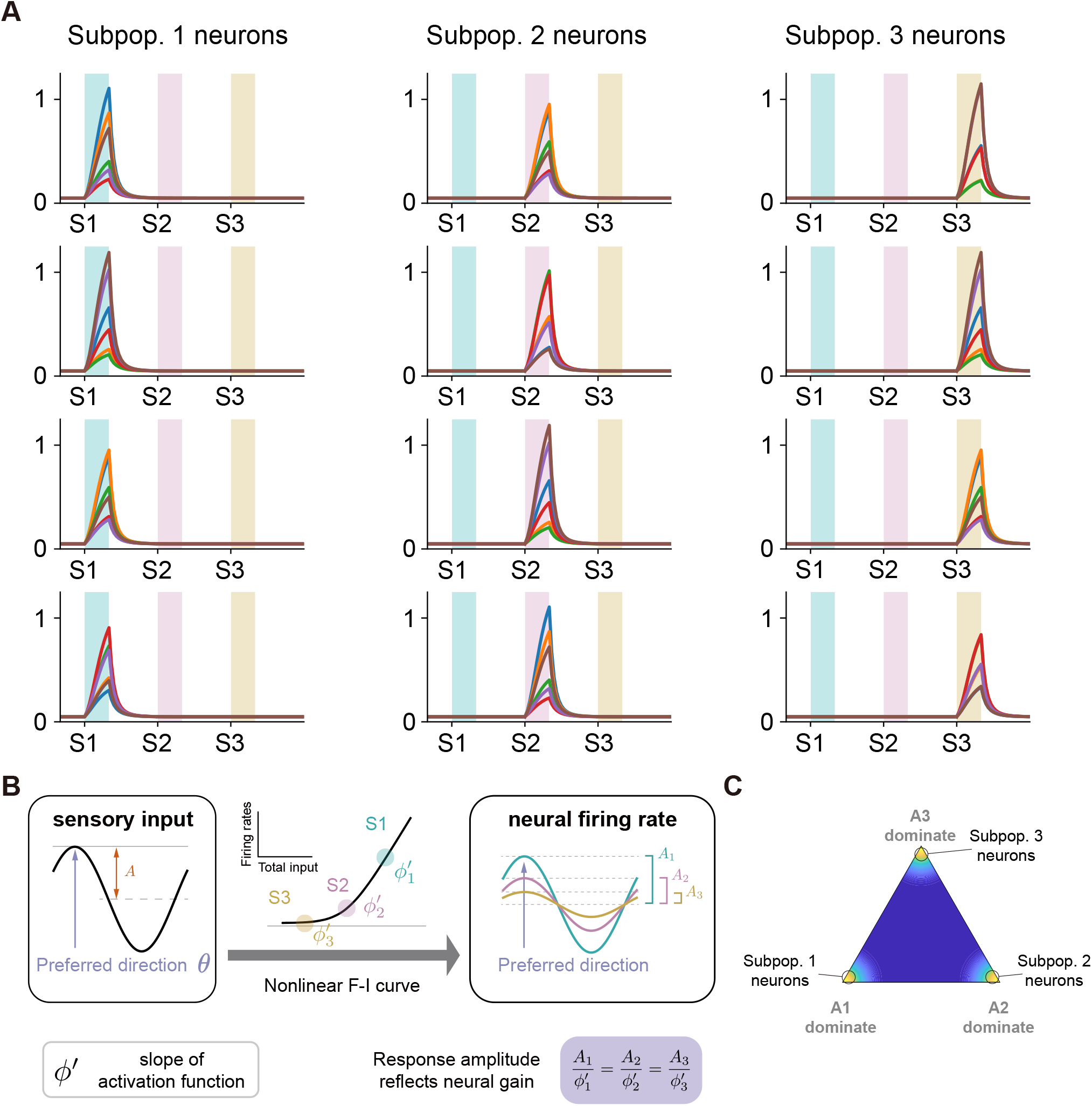
A naive hypothesis of the subpopulation activity for the SWM task. (A) Example neurons of the three subpopulations in the alternative hypothesis, where different subpopulations activate during distinct input periods (e.g., subpopulation 1 during S1, subpopulation 2 during S2, etc.). (B) To measure a neuron’s tuning to the six items, we first compute the conditional average firing rates and fit them with a cosine function. This gives us two quantities: the amplitude *A* and the preferred direction *θ*. The amplitude *A* reflect the sensitivity of the neuron to stimulus. (C) Distribution of neuronal tuning amplitude. The neurons in the three subpopulations of panel A distribute at distinct apexes.

**Fig. S4-4.**
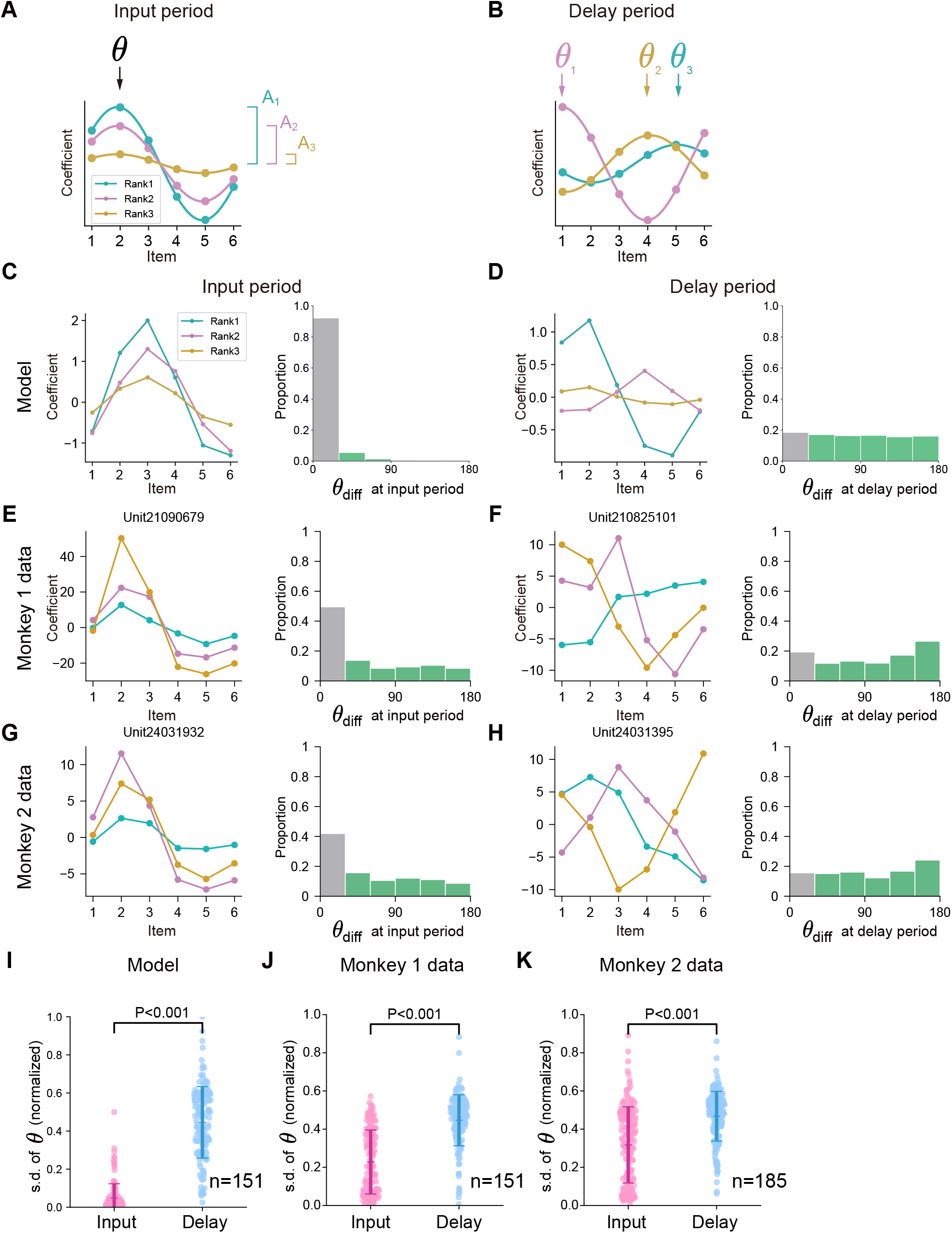
Example neurons and the neuronal tuning patterns of the Restricted-RNN model and the experiment data for the SWM task. (A and B) We use cosine function to fit each neuron’s tuning curve. In this way, the tuning pattern is captured by the phase **ϕ** and amplitude *A* of the resulting cosine curve. Item-selective neurons in the input period share similar tuning phase across ranks. While during the delay period, the tuning phases across ranks of the item-selective neurons therein are uniformly distributed. (C, E, and G) Neuronal tuning patterns of model (C) and monkey data (E and G) during the input period. (left) Tuning curves of example neurons with respect to the six locations. (right) Distribution of tuning phase difference across ranks. (D, F, and H) Same as (C, E, and G) but for the Delay period. (I) Comparison of the standard deviations of phase different distribution in the input and delay periods in our model. The phase difference in the input period is significantly smaller than the one in the delay period. (J and K) Same as (I) but for Monkey 1 (J) and Monkey 2 (K), respectively. In both cases, the phase difference in the input period is also significantly smaller than the one in the delay period.

**Fig S4-5.**
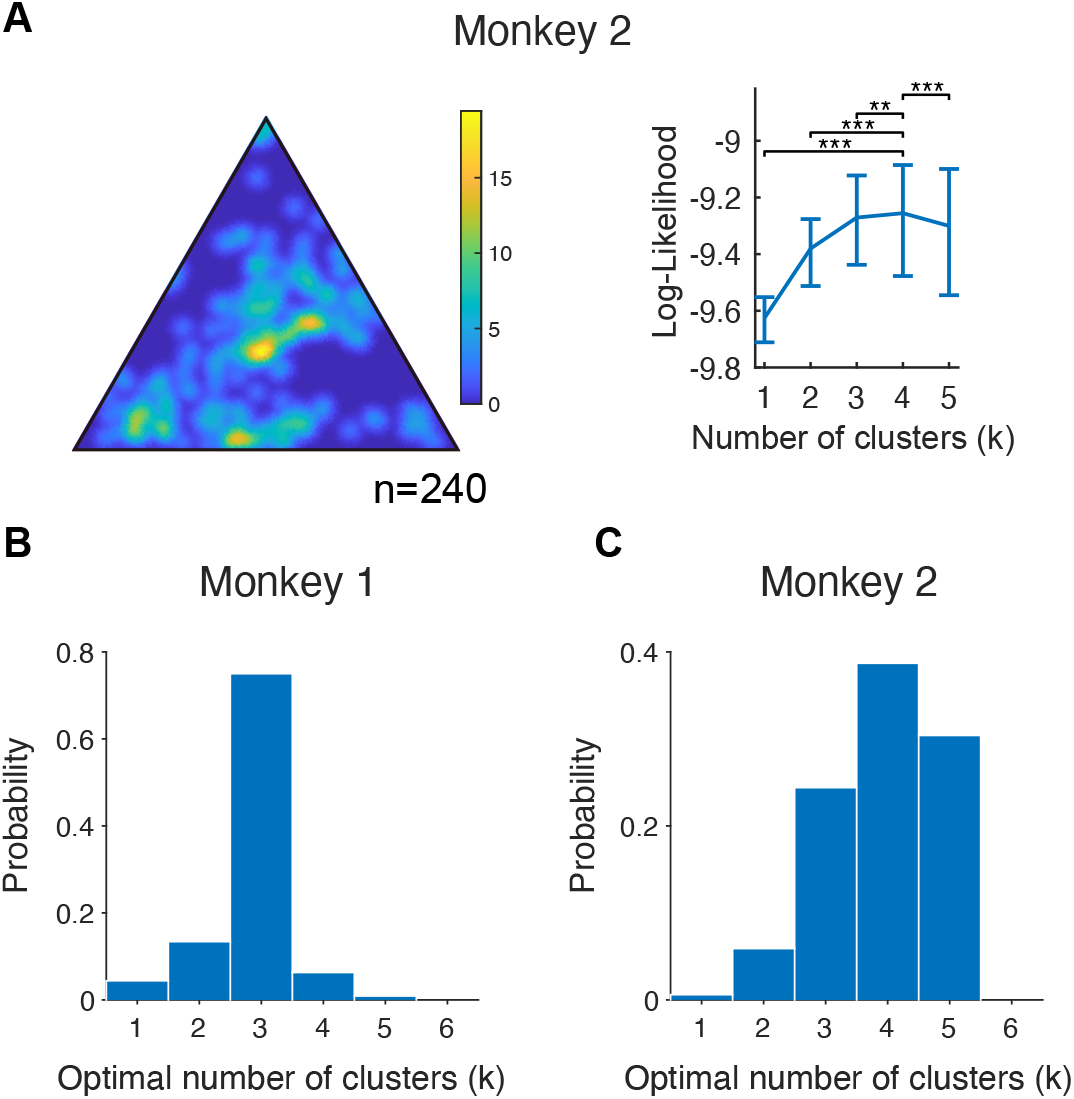
Clustering analysis of monkey data in the SWM task. (A) Distribution of neuronal tuning amplitudes for Monkey 2. Clustering analysis suggests four clusters (bootstrapped; median log-likelihood with IQR error bars; one-side Wilcoxon signed-rank test with Benjamini-Hochberg FDR correction for multiple comparisons ***p<0.001). (B and C) Distribution of optimal number of clusters (k) among 100 bootstraps for Monkey 1 (B) and Monkey 2 (C).

**Fig S4-6.**
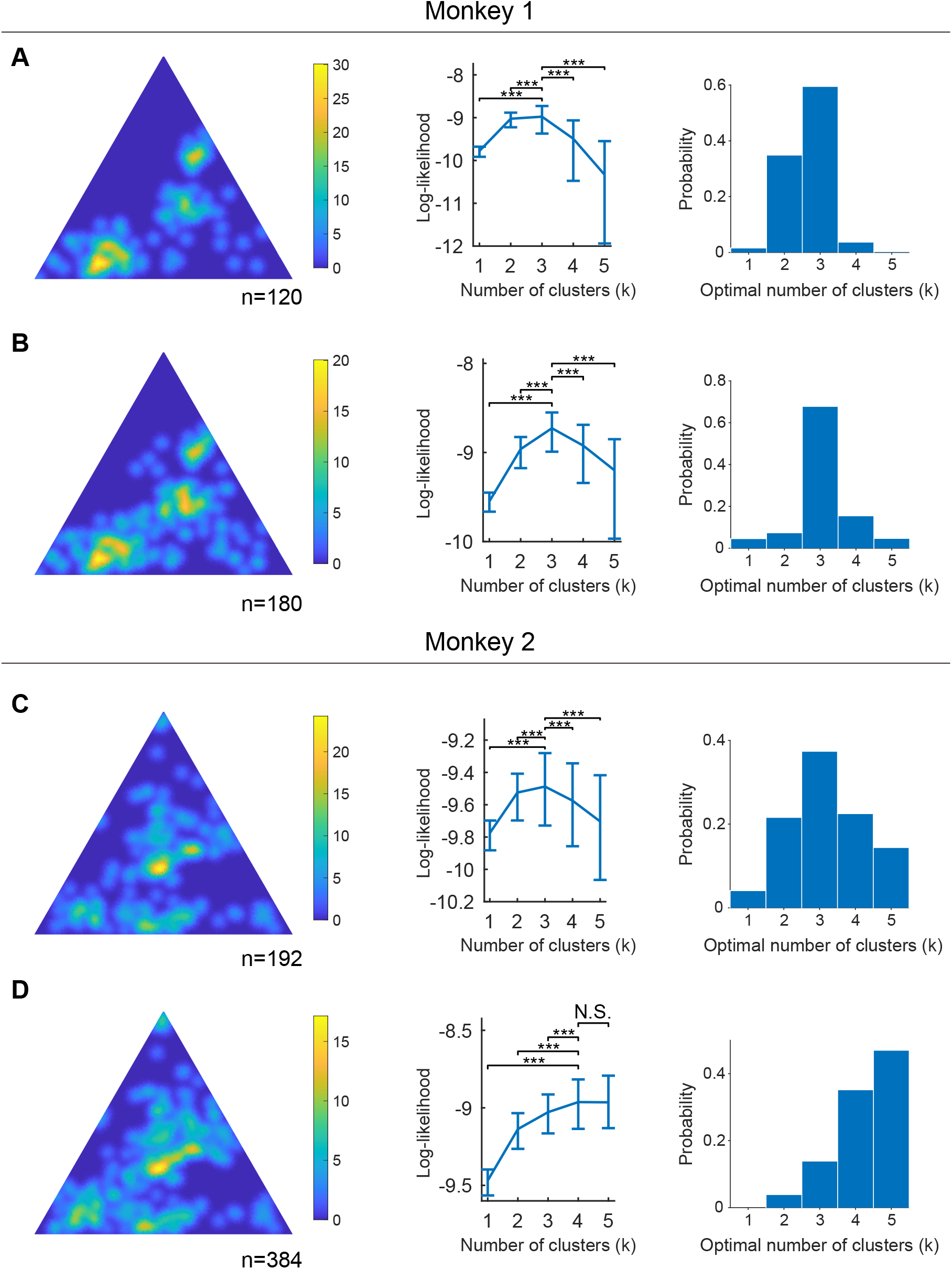
Clustering analysis of both monkeys in the SWM task using various thresholds. Clustering analysis (see Methods) with a lower and a higher threshold than the one used in the main text for choosing item-selective neurons during the input period. Results show the median log-likelihood with IQR error bars across different number of clusters (bootstrapped; one-side Wilcoxon signed-rank test with Benjamini-Hochberg FDR correction for multiple comparisons ***p<0.001 and N.S. >=0.05). (A and B) Clustering analysis suggests three clusters in the distribution of neuronal tuning amplitudes for Monkey 1. (C) With lower threshold, clustering analysis suggests three clusters in the distribution of neuronal tuning amplitudes for Monkey 2. (D) With higher threshold, clustering analysis suggests four or five clusters in the distribution of neuronal tuning amplitudes for Monkey 2. In this situation, more clusters fit the distribution better.

**Fig. S5-1.**
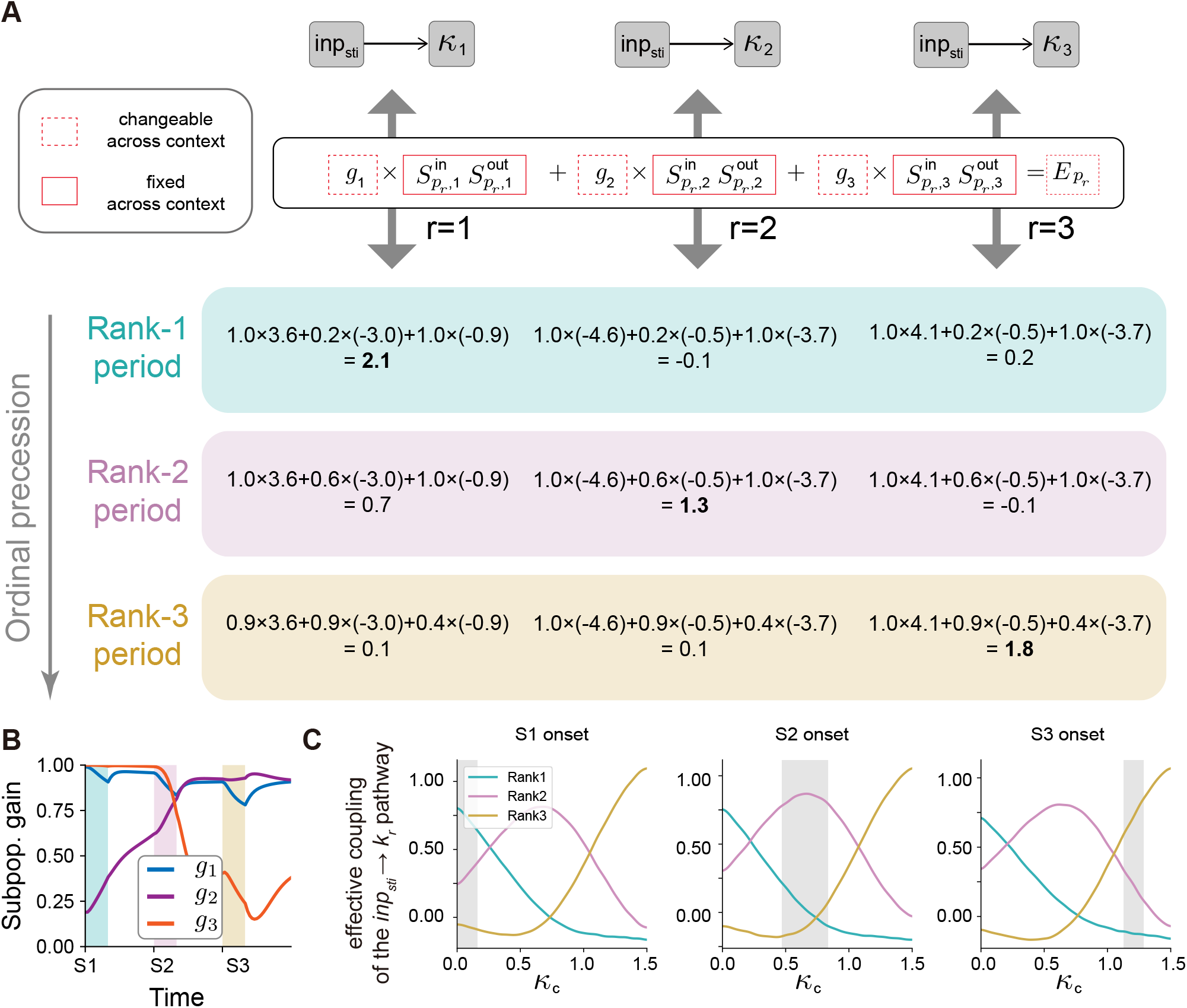
Mean-field analysis of the Restricted-RNN for the SWM task. (A) Mean-field analysis. An example of computing effective coupling during the sampling period S1, S2, and S3. (B) Subpopulation gain curves during the sampling period. (C) Temporal profile of the effective coupling of the *inp*_sti_ → *k*_*r*_ pathway as a function of the value of factor *k*_*c*_. During the *r*-th stimulus onset, the factor *k*_*c*_ changes in a way such that the effective coupling strength of the *inp*_sti_ → *k*_*r*_ pathway became highest (i.e., opening the *inp*_sti_ → *k*_*r*_ pathway and closing other pathways).

**Fig. S5-2.**
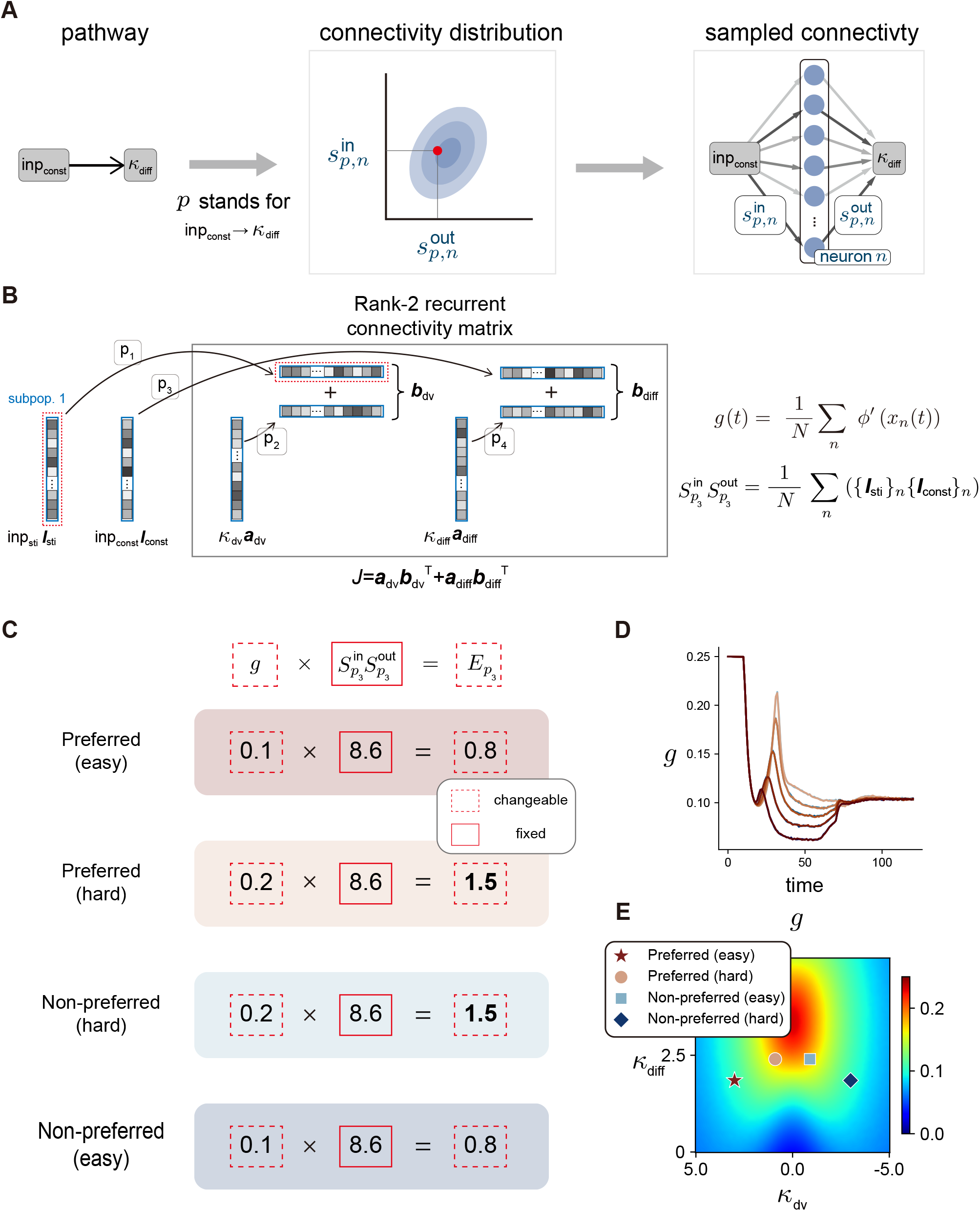
Mean-field analysis of the Restricted-RNN for the PDM task. (A) Pathway-based generative model for the *inp*_const_ → *k*_diff_ pathways. There is only one subpopulation in this Restricted-RNN model (see Fig. S2-7 for a general description). (B) Computation of the collective level variables in the implementational level low-rank RNN. (C) Mean-field analysis. An example of computing effective coupling for stimulus of different coherences (easy and hard) under preferred and non-preferred contexts. (D) Temporal profile of the population gain for subpopulation 1 (i.e., *g*) with different absolute stimulus strengths. (E) Heat map of *g* in the *k*_dv_-*k*_diff_ space. Input states of the same coherence possess different gain regions under different contexts. For example, the easy state under preferred context has lower *g* compared to the one of easy state under non-preferred context.

**Fig. S5-3.**
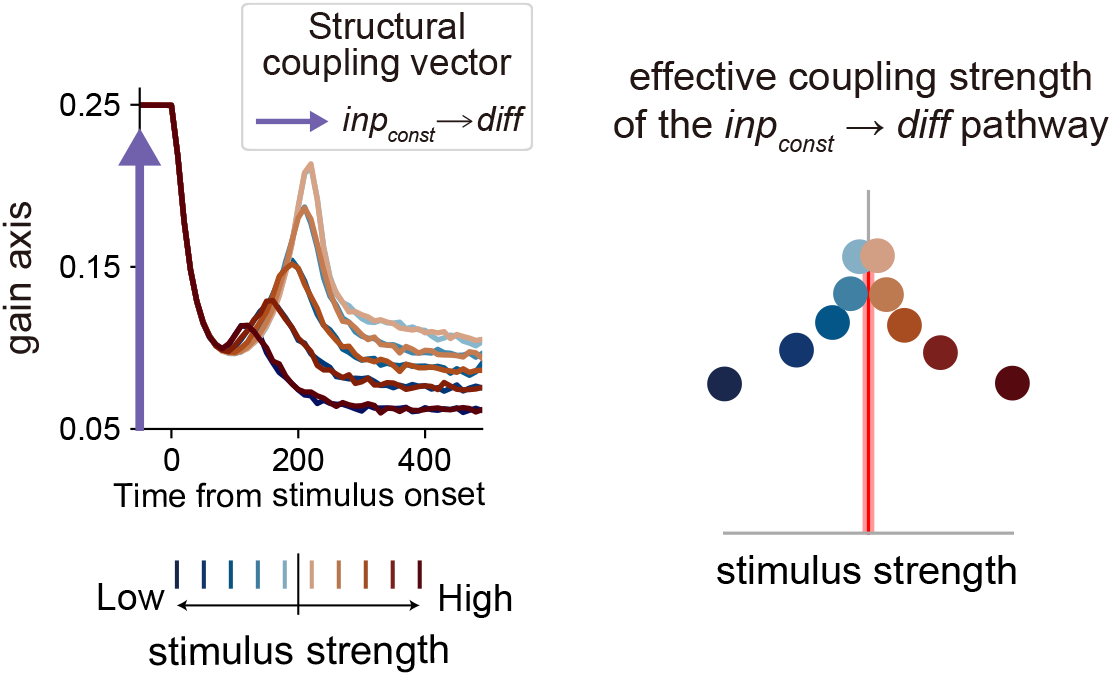
Mechanism behind difficulty representation in the PDM model. Despite there is no direct difficulty input, the subpopulation gain evolves differently with different absolute stimulus strengths. This will modulate how factor *k*_*diff*_ integrates the stimulus-irrelevant input and generates difficulty representation.

**Fig. S5-4.**
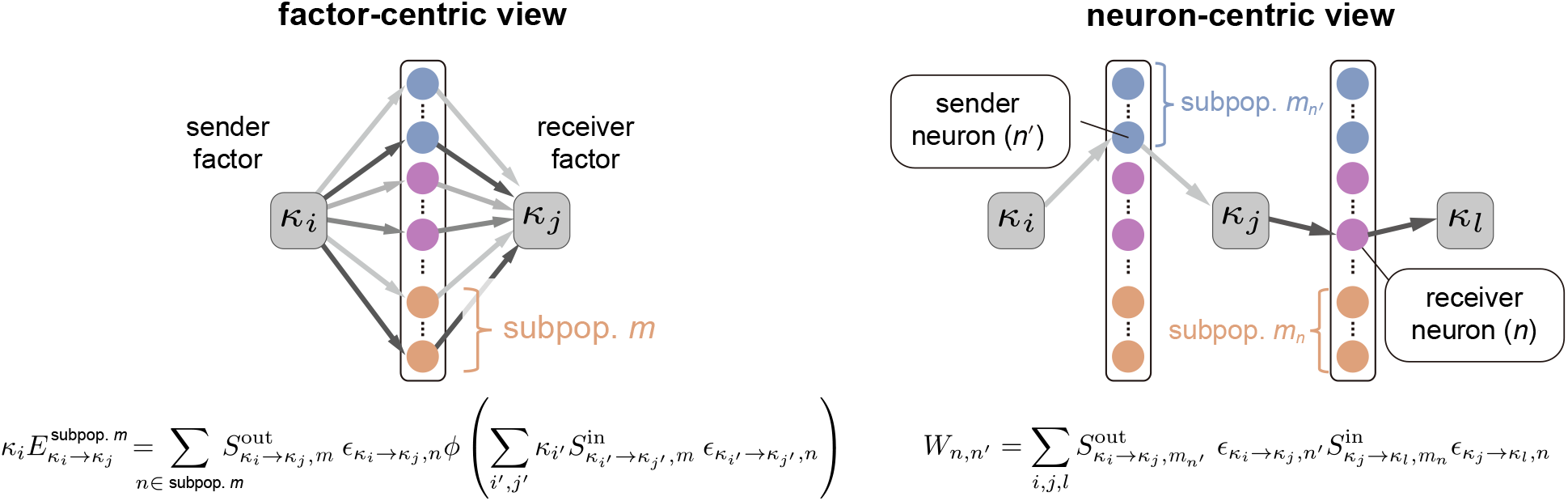
Key ideas underlying the construction of the implementational-level Restricted-RNN. (A) Comparison between the factor-centric view (left) and the neuron-centric view (right) for Restricted-RNN. The factor-centric view better describes communication among factors, while the neuron-centric view is more appropriate for building the implementational-level low-rank RNN.

## Supplementary Note

### Supplementary Note 1 (Mean-field theory of low-rank RNNs)

Understanding how a network’s connectivity relates to its function is a fundamental challenge in the study of recurrent neural networks (RNNs). This problem is particularly complex for general RNNs due to their high-dimensional and nonlinear dynamics. Recently, a theoretical framework based on mean-field theory has been introduced (Sompolinsky et al., 1988; Beiran et al., 2021; Dubreuil et al., 2022), demonstrating that the dynamics of low-rank networks can be expressed in an explicit form when the connectivity strength of each neuron is randomly sampled from a multivariate Gaussian mixture model.

Consider a low-rank RNN with *R* ranks and *N* neurons, where the dynamics are described by the following equation:

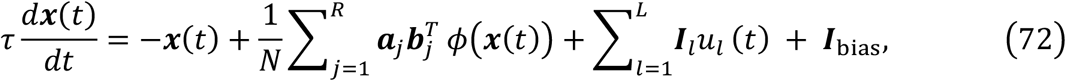

where *τ* is the time-constant, ***x***(*t*) is the *N*-dimensional hidden state at time *t, ϕ* is a non-linear activation function, and ***a***_*j*_ and ***b***_*j*_ are *N*-dimensional vectors representing the *j*-th left and right connectivity vectors of the connectivity matrix, respectively. The term ***I***_*l*_ represents the *l*-th input connectivity vector, and *u*_*l*_(*t*) is the corresponding time-dependent input signal. The constant vector ***I***_bias_ is an additive bias term. The network’s output is defined as: 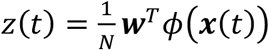, where ***w*** ∈ *R*^*N*^ is the readout vector. Without loss of generality, the bias term can be treated as an input with a constant drive *u*_bias_ ≡ 1. Therefore, in rest of the section, we omit the biased term.

In network (1), its hidden state ***x***(*t*) is constrained in the subspace spanned by the left connectivity vectors {***a***_*j*_} and input connectivity vectors {***I***_*l*_}, and can be expressed as:

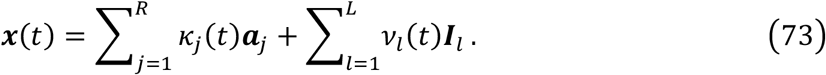

where *k*_*j*_(*t*) and *ν*_*l*_(*t*) are the task and input factors associated with ***a***_*j*_ and ***I***_*l*_, respectively. Together, these factors are referred to as latent factors. The dynamics of these factors are termed as latent dynamics, given by:

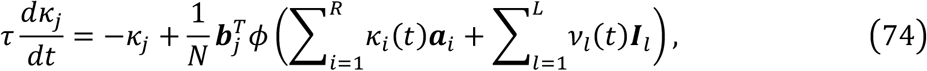

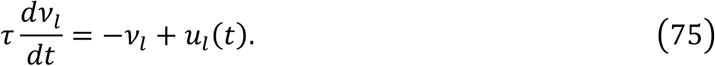

Note that each neuron here is described by its 2*R* + *L* components on the connecticity vectors ***a***_*j*_ and ***b***_*j*_ and input connectivity vectors ***I***_*l*_. Assume that the 2*R* + *L* components are sampled from a multivariate Gaussian mixture model. That is, the joint probability density of (*I*_1,*n*_, …, *I*_*L,n*_, *a*_1,*n*_, …, *a*_*R,n*_, *b*_1,*n*_, …, *b*_*R,n*_) is given by:

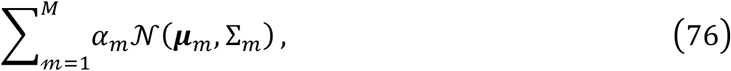

where *M* is the number of clusters (termed as subpopulations here), *α*_*m*_ is the weight of the *m*-th subpopulation, and each subpopulation is a (2*R* + *L*)-dimensional Gaussian distribution with mean ***μ***_*m*_ and covariance matrix Σ_*m*_. Under these assumptions, in the mean-field limit (*N* → ∞), the dynamics of the task variables can be described by:

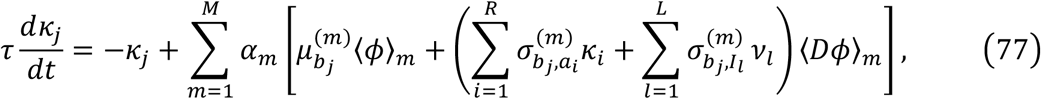

where 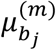 is the mean of *b*_*j*_ in the *m*-th subpopulation, and 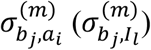 is the covariance of *b*_*j*_ with *a*_*i*_ (*I*_*l*_ ) in the *m*-th subpopulation, as defined in Σ_*m*_ . The term ⟨*ϕ*⟩_*m*_ and ⟨*Dϕ*⟩_*m*_ are the average activity and average **gain** of the *m*-th subpopulation, respectively, and are defined as:

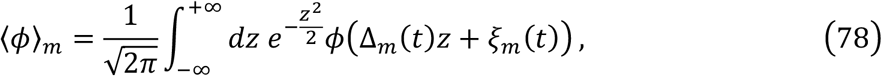

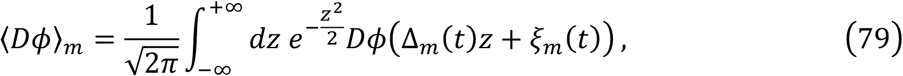

where 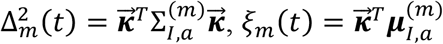. Here 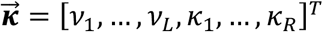 is the vector of all input and task factors, 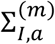 is the submatrix of Σ_*p*_ corresponding to the set of variable {*I*_1_, …, *I*_*L*_, *a*_1_, …, *a*_*R*_}. Similarly, 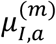 is the subset of the mean vector ***μ***_*m*_ that corresponds to these variables. For simplicity, in rest of the paper, we further assume 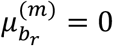 for all ranks *r* and subpopulations *m*, reducing the dynamics of task variables to:

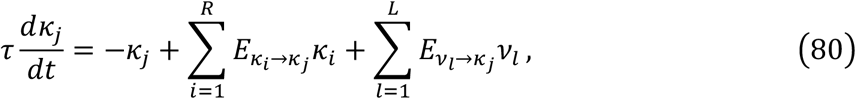

where 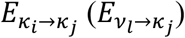 is the effective coupling strength of the *k*_*i*_ (*ν*_*l*_) to *k*_*j*_ pathway defined by:

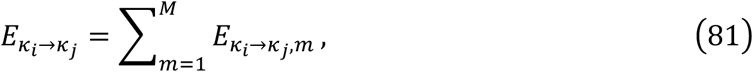

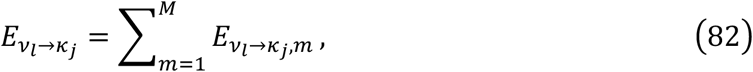

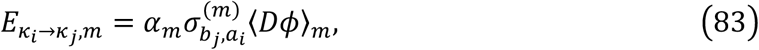

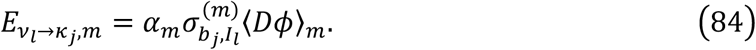

Here each 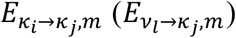 stands for the effective coupling strength mediated by the *m*-th subpopulation.

#### Necessity of multiple subpopulations

The capacity of a network with only one single subpopulation is limited. Consider a network containing two input variables *ν*_1_ and *ν*_2_ and one decision variable *k*_*dv*_ is designed to handle context-dependent computation. In context 1, *k*_*dv*_ should receive the information from *ν*_1_, whereas in context 2, it should instead receive input from *ν*_2_. If the network only has one subpopulation, the effective coupling strength of *ν*_1_ → *k*_d*v*_ and *ν*_2_ → *k*_*dv*_ are both controlled by the same value ⟨*Dϕ*⟩_1_, which means that the two paths are either simultaneously active or inactive. This constraint (one subpopulation) prevents the network from selectively routing different inputs in different contexts. As a conclusion, incorporating multiple subpopulations enables the network to support more flexible computations (Dubreuil et al., 2022).

### Supplementary Note 2 (the derivation of Restricted-RNN dynamics associated with factor graph)

For Restricted-RNN, let ***x***(*t*) is the *MN*-dimensional activation vector and the activity vector is given by ***r***(*t*) = *ϕ*A***x***(*t*)B . The terms 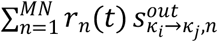 and 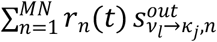 capture the information flowing from *k*_*i*_ to *k*_*j*_ and from *ν*_*l*_ to *k*_*j*_, respectively. Summing over all sender variables yields the total input to *k*_*r*_:

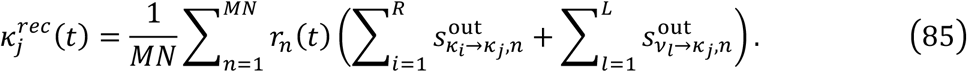

Next, the total input to unit *n* is as following:

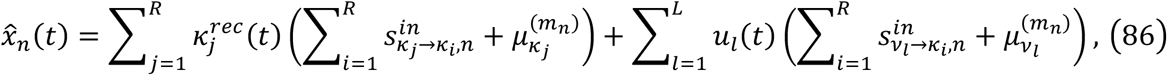

where 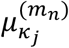 and 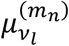 are parameters representing the subpopulation specific mean value for the corresponding task variable. At the implementational level, the RNN dynamics follow:

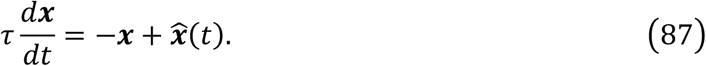

It can be verified that the instantiated RNN is a rank-*R* low-rank network described by:

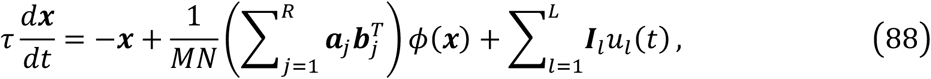

where ***a***_*j*_ is a (*MN*)-dimensional vector with element

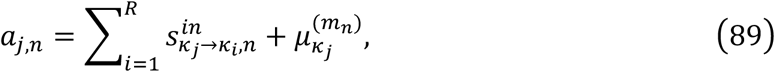

***b***_*j*_ is a (*MN*)-dimensional vector with element

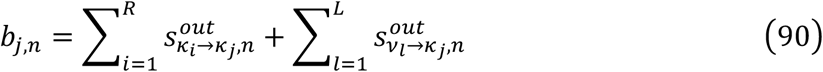

and ***I***_*l*_ is a *MN*-dimensional input vector with element

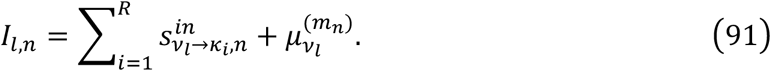

### Supplementary Note 3 (the derivation of factor dynamics in the mean-field limit)

The hidden state can be decomposed in the following way:

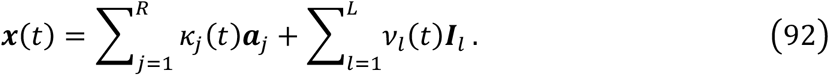

By plugging this decomposition into the RNN dynamics, we obtained

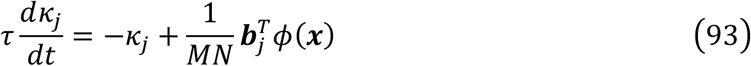

Note that

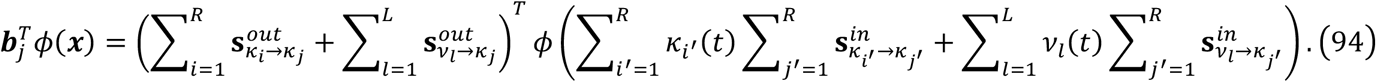

To compute 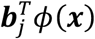, let us first consider

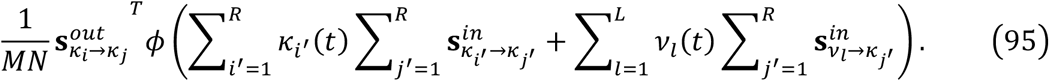

In the mean-field limit (*N* → ∞), by applying Stein’s lemma, this term is equal to

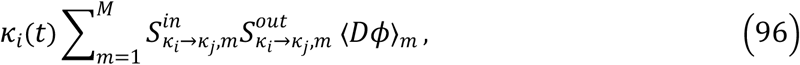

where ⟨*Dϕ*⟩_*m*_ is the average gain of the *m*-th subpopulation. This result demonstrated that the output connectivity vector can read out the information conveyed by the same pathway and is inaccessible to the information conveyed by a different pathway. For brevity, we denote this mean-field value as 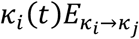.

Following the similar derivation, we can compute the mean-field value of

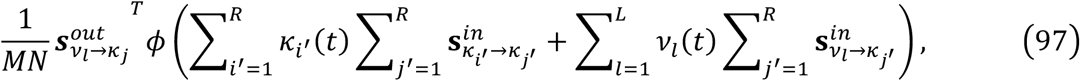

which is equal to

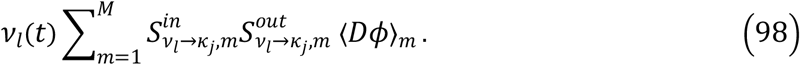

Similarly, we denote this value as 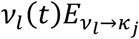.

Taken together, we complete the derivation of the factor dynamics in the mean-field limit:

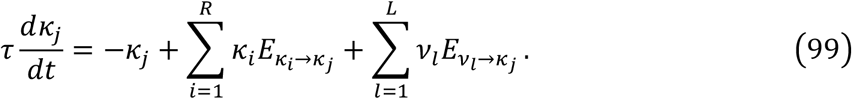

### kiSupplementary Note 4 (the universal approximation property for the Restricted-RNN)

In this section, we formalize a universal approximation property for the Restricted-RNN at the level of collective dynamics. We show that, under mild assumptions on the activation nonlinearity and with an appropriate parameterization, the collective-variable dynamics induced by a Restricted-RNN can approximate any target vector field on a compact set to arbitrary precision.

We consider collective dynamics in ℝ^*R*^ of the form,

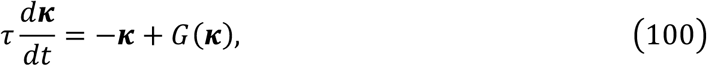

where G: ℝ^*R*^ → ℝ^*R*^ is a continuous vector field, and *τ* is a time constant. Let *K* ⊂ ℝ^*R*^ be a compact set. Our goal is: for any *ϵ* > 0, construct a Restricted-RNN with parameter set Θ^∗^ that the induced mean-field dynamics

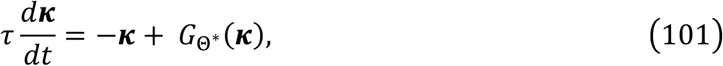

satisfy

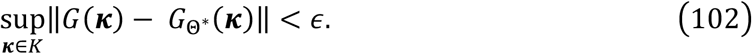

For this goal, we construct a Restricted-RNN with *M* subpopulations, *R* latent factor and a biased term. There is no direct latent factor to latent factor pathways, meaning that for any *i, j* = 1, …, *R* and subpopulation index *m*,

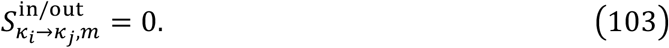

There is a pathway from the bias factor to the latent factor *k*_*i*_, parameterized as

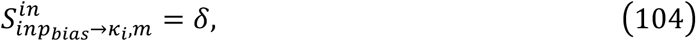

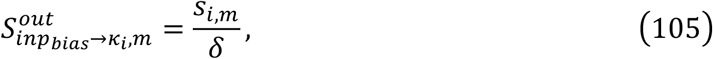

with *δ* > 0 and *s*_*i,m*_ are free parameters. Let ***s***_*m*_ = [*s*_1,*m*_, …, *s*_*R,m*_] and 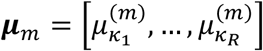.

Under this construction, the mean-field dynamics of the latent factors take the form

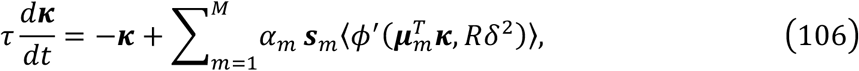

where 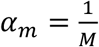 is the subpopulation weight. The free parameters in this network are

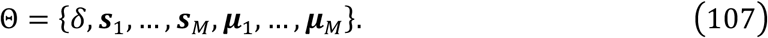

Define a function *F* on the set *K* × [0, 1] by:

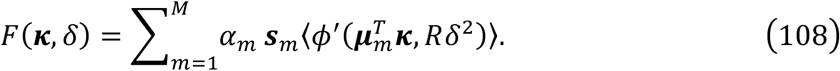

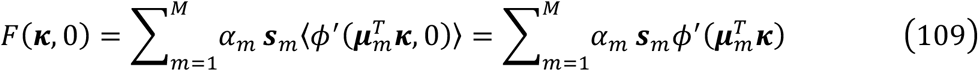

We assume *ϕ*′ is nonpolynomial and continuous. Under these conditions, by the universal approximation theorem, there exists a finite *M* and parameters 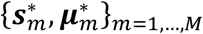 such that

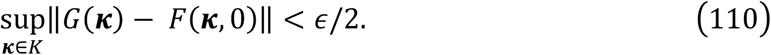

For these fixed 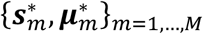, the mapping (***κ***, *δ*) → *F*(***κ***, *δ*) is continuous on *K* × [0, 1].

Since *K* × [0,1] is compact, this continuity is uniform. Hence, there exists *δ*^∗^ ∈ (0,1] such that,

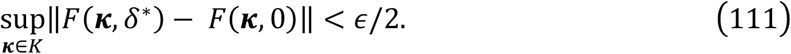

Combining these two steps yields

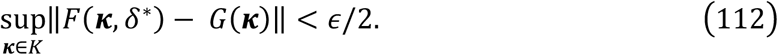

This establishes that the Restricted-RNN can approximate the target vector filed on *K* to arbitrary precision.

